# A Spike Trimer Dimer-Inducing Nanobody with Anti-Sarbecovirus Activity

**DOI:** 10.1101/2024.06.19.598823

**Authors:** Iris C. Swart, Oliver J. Debski-Antoniak, Aneta Zegar, Thijs de Bouter, Marianthi Chatziandreou, Max van den Berg, Ieva Drulyte, Krzysztof Pyrć, Cornelis A.M. de Haan, Daniel L. Hurdiss, Berend-Jan Bosch, Sabrina Oliveira

## Abstract

The continued emergence and zoonotic threat posed by coronaviruses highlight the urgent need for effective antiviral strategies with broad reactivity to counter new emerging strains. Nanobodies (or single-domain antibodies) are promising alternatives to traditional monoclonal antibodies, due to their small size, cost-effectiveness and ease of bioengineering. Here, we describe 7F, a llama-derived nanobody, targeting the spike receptor binding domain of sarbecoviruses and SARS-like coronaviruses. 7F demonstrates potent neutralization against SARS-CoV-2 and cross-neutralizing activity against SARS-CoV and SARS-like CoV WIV16 pseudoviruses. Structural analysis reveals 7F’s ability to induce the formation of spike trimer dimers by engaging with two SARS-CoV-2 spike RBDs, targeting the highly conserved class IV region. Bivalent 7F constructs substantially enhance neutralization potency and breadth, up to more recent SARS-CoV-2 variants of concern. Furthermore, we demonstrate the therapeutic potential of 7F against SARS-CoV-2 in the fully differentiated 3D tissue cultures mirroring the epithelium of the human airway ex vivo. The broad sarbecovirus activity and distinctive structural features of 7F underscore its potential as promising antiviral against emerging and evolving sarbecoviruses.

## Introduction

In recent years, two severe acute respiratory syndrome (SARS)-related coronaviruses (SARS-CoVs) have crossed over from zoonotic reservoirs to humans, causing the SARS-CoV epidemic in 2003 and the SARS-CoV-2 pandemic declared in 2020 (1,2). Both SARS-CoV-2 and SARS-CoV belong to the subgenus *Sarbecovirus* within the genus *Betacoronavirus*, which otherwise comprises a large variety of SARS-like coronaviruses that circulate mainly in bats, including high-risk strains such as RaTG13, WIV1, SARS-like CoV WIV16 and RsSHC014 (2–4). Given the potential for new zoonotic spillover events of viruses within this subgenus, there is an urgent need for broad-spectrum therapeutics capable of targeting conserved epitopes across the subgenus *Sarbecovirus*. SARS-CoV-2 and SARS-CoV utilize the angiotensin-converting enzyme 2 (ACE2) as their entry receptor, which they engage via the receptor-binding domain (RBD) present within the spike (S) glycoprotein (2,5,6). In its prefusion state, the spike protein forms a homotrimer and undergoes significant structural changes to regulate the exposure and availability of the RBD. This occurs through an alternating “up” and “down” mechanism, wherein the RBD transitions between a state that is accessible to receptors (in the “up” conformation) and a state that is not accessible to receptors (in the “down” conformation) (7,8). The RBD has become a main target for therapeutic development. During the SARS-CoV-2 pandemic, research on nanobodies gained momentum alongside traditional monoclonal antibody (mAb) approaches (9).

Nanobodies, also known as single-domain antibodies (sdAbs) or VHHs (Variable Heavy domain of Heavy chain antibodies), constitute the variable domain of heavy-chain antibodies found in, amongst others, camelids and cartilaginous fishes (10–12). These small binding fragments of approximately 15 kDa, are extremely stable, can facilitate excellent tissue penetration and can easily be bio-engineered into multivalent formats (13–15). In contrast to conventional antibodies, nanobodies can be produced at low-cost in bacterial systems and their high stability and small size allows for intranasal administration (16). For respiratory pathogens, like sarbecoviruses, intranasal delivery is preferred for localized high concentrations, faster onset, and reduced systemic exposure (17). These attributes combined make nanobodies highly valuable for engineering innovative biotherapeutics with potent and broad antiviral activity against viral pathogens.

Targeting conserved epitopes on the CoV S, particularly within the RBD, offers multiple advantages: broad coverage across various sarbecovirus strains, heightened resistance to neutralization escape, and the potential to counteract future emerging strains. Notably, several nanobodies with broad sarbecovirus activity have been described, of which the majority targets the RBD (18–25). Two conserved epitopes on the spike RBD have been well characterized, the class III and IV epitopes, as classified by Barnes et al., (26–28). Multiple class IV antibodies, broadly targeting sarbecoviruses have been identified and characterized during the pandemic (29–33). The occluded nature of the class IV epitope likely contributes to its conserved characteristics amongst the sarbecoviruses.

Here, we present a novel class IV epitope nanobody, 7F, with broad neutralization activity towards sarbecoviruses. We structurally characterized the spike binding mode of this nanobody and evaluated its neutralization potency and breadth. The binding of 7F allows simultaneous interaction with the RBDs of two spike proteins, inducing S-ectodomain trimer dimers - a phenomenon which has only been reported previously twice (24,34). Additionally, we evaluated the neutralization potency and breadth of 7F after dimerization. The combination of targeting a conserved epitope and displaying broad reactivity towards sarbecoviruses, positions 7F as a promising candidate for combating future outbreaks.

## Results

### Identification of broadly reactive sarbecovirus targeting nanobody 7F

To elicit an immune response and isolate sarbecovirus S-targeting nanobodies, two llamas were immunized with prefusion trimeric spike proteins of multiple betacoronaviruses, including SARS-CoV and SARS-CoV-2. Thereafter, blood was collected and phagemid libraries were generated and pooled for the nanobody selection process. Two consecutive rounds of phage panning against SARS-CoV-2 RBD, followed by phage ELISAs, resulted in five unique sequences of promising nanobody candidates. Further characterization of the purified monovalent nanobodies revealed their high-affinity binding to the SARS-CoV-2 spike protein and their capacity to neutralize SARS-CoV-2 pseudovirus (**Fig. S1A-B**). Given the study’s focus on developing broad neutralizing sarbecovirus nanobodies, we evaluated their cross-reactivity to SARS-CoV and SARS-like CoV WIV16. Notably, two of the selected nanobodies, namely nanobody 7F and 9A, demonstrated binding to SARS-CoV and SARS-like CoV WIV16 as well (**Fig. S1C**). Furthermore, both nanobodies were capable of cross-neutralization of SARS-like CoV WIV16 pseudovirus (**Fig. S1D**).

Of the two cross-reactive nanobody candidates, 7F was selected for further analysis, as it exhibited low nanomolar (nM) binding affinity to all three sarbecoviruses in ELISA (**Fig. 1A**) and demonstrated broad neutralization activity against pseudoviruses of SARS-CoV-2, SARS-CoV and SARS-like CoV WIV16, with IC_50_ values falling in the nM range (33.4 ± 17.2, 27 ± 5 and 4.7 ± 0.9 nM respectively; **Fig. 1B**). To evaluate the epitope location of 7F on the spike protein, biolayer interferometry (BLI) was performed. The strong binding signal of 7F to the spike RBD in contrast to the spike protein N-terminal domain (NTD) reveals that 7F binds to an epitope on the RBD (**Fig. 1C-D**).

**Figure 1.**
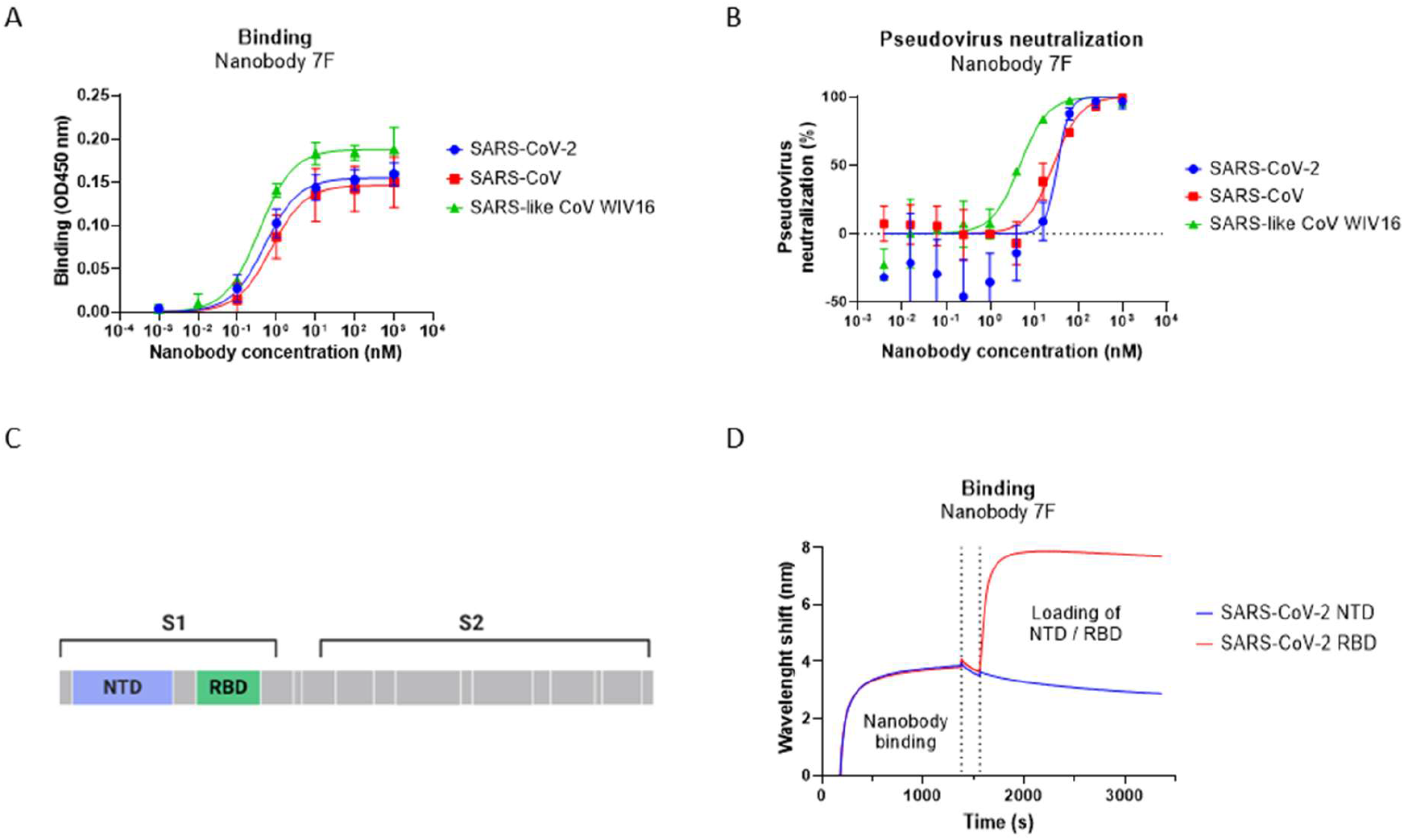
Camelid-derived nanobody 7F shows cross-reactivity activity against sarbecoviruses. **A.** ELISA-based reactivity of 7F to plate-immobilized spike ectodomain of SARS-CoV-2, SARS-CoV and RBD of SARS-like-CoV WIV16. Data points represent the mean ± SDM of n = 3 replicates from one representative of three independent experiments. **B.** Neutralization of SARS-CoV-2, SARS-CoV and SARS-like CoV WIV16 pseudovirus by serial diluted nanobody on VeroE6 cells. Data points represent the mean ± SDM of n = 3 replicates from one representative of two independent experiments. **C.** Schematic representation of the SARS-CoV-2 spike protein, with NTD and RBD labeled **D.** Binding of 7F to different spike domains analyzed by BLI. 7F was immobilized on NTA sensors, after which 7F was saturated in binding with either SARS-CoV-2 NTD or RBD. Data shown is based on a single experiment.

### 7F induces formation of SARS-CoV-2 spike trimer dimers

To gain insight into the interaction site of 7F, cryo-electron microscopy (cryo-EM) single particle analysis was performed on the prefusion-stabilized SARS-CoV-2 ancestral Wuhan spike ectodomain trimer complexed with 7F (**Fig.2**). Two-dimensional classification analysis revealed only particles representing a head-to head spike trimer dimer (**Fig.S2A**). Three-dimensional classification further confirmed this phenomenon, with a single class representing a trimer dimer observed. All six RBDs are in the “up” conformation and subsequently interact with six molecules of 7F (**Fig.2A-C**). Further processing yielded a final reconstruction with a global resolution of 3.3 Å (**Fig.S2-3**). Local refinement improved the resolution at the interface to 3.1 Å (**Fig.2A + B and Fig.S2-3**), allowing accurate modelling and characterization of the interacting residues at the 7F-RBD interfaces.

**Figure 2.**
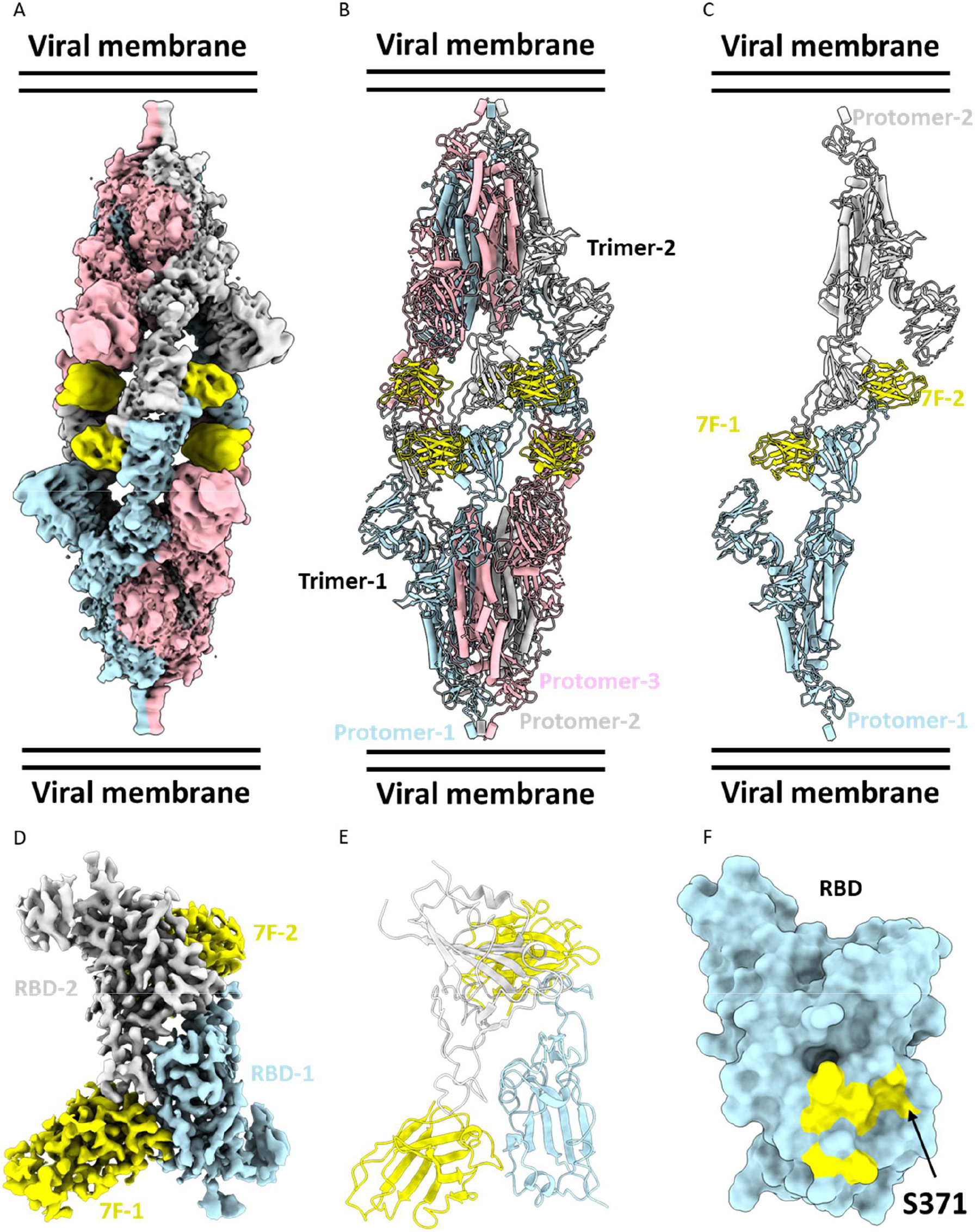
Structural analysis of 7F in complex with the SARS-CoV-2 spike trimer. **A**. EM density map. The complex comprises six 7F molecules bound to two SARS-CoV-2 spike trimeric ectodomains. **B.** Atomic model of the six 7F molecules bound to two SARS-CoV-2 spike trimeric ectodomains. Spike protein protomers are in blue, grey, and pink, respectively. The 7F molecules are in yellow. **C.** Atomic model of a single protomer of each spike trimer, bound to two 7F molecules. Protomer-1 is blue, protomer-2 is grey and both 7F molecules are represented in yellow. **D.** Locally refined EM density map of the interaction site between two 7F molecules and two SARS-CoV-2 RBDs. **E.** Atomic model of the interaction site. RBD-1 is blue, RBD-2 is grey and 7F molecules are represented in yellow. **F.** Surface representation atomic model of SARS-CoV-2 RBD (blue) with the 7F epitope highlighted (yellow).

The 7F-RBD complex contains three binding interfaces. Two of these interfaces are between the juxtaposed RBDs and a 7F molecule, labelled as ‘interface-major’ and ‘interface-minor’ based on the surface area buried in each RBD-7F interaction. While the third interface is between opposing RBDs and labelled “RBD-RBD-interface”. Interface–major constitutes a buried surface area of 638 Å^2^ and is likely solely responsible for the single dominant binding affinity measured in ELISA assays (**Fig.1A**). The interface-minor consists of a buried surface area of 165 Å^2^, predominantly involving residues in the CDR3 loop, but also a single residue from framework-2 of 7F. Surprisingly, the proximity in which the interaction between RBDs and 7Fs in the dimer formation takes place, allowing for a third interaction between both RBDs. This “RBD-RBD” interface creates a further buried surface area of 1008 Å^2^ between the receptor binding motif of each RBD.

The interface-major interaction coincides with that of class IV anti-RBD mAbs (**Fig. 2C**). To understand this major interaction, we compared a range of both antibodies and nanobodies from class IV (**Fig. S4**). Monoclonal antibodies such as S2X259 (30), EY6A (32), CR-3022 (29), S304 (31) and nanobody VHH72 (25) largely overlap the binding surface of 7F, while the monoclonal antibodies H014 (33) and S2A4 (31) encompass the entirety of the 7F interface-major binding surface. These antibodies and nanobodies exclusively bind SARS-CoV-2 RBD in an “up” conformation, but do not induce trimer dimer formation. It is likely that the interactions observed here at site Interface-minor drive the dimerization resulting in the RBD-RBD interface, through shape-complementation.

The antibody cluster S2A4, EY6A, CR3022 and S304 have been shown to be capable of triggering S1 shedding by Cryo-EM (31,32,35). The binding sites for these antibodies are completely buried in the spike ectodomain when RBD is in the “down” conformation, with at least a “2-up” RBD conformation required for interaction, due to clashes of the Fab fragments with the adjacent “down” RBD, S2 subunit and NTDs (32,35). In the context of the closed S, the 7F binding site is occluded. Superimposition of 7F to “1-up” and “2-up” SARS-CoV-2 S-ectodomains, suggest only a “3-up” conformation would allow binding (**Fig. S5**), due to clashes with adjacent RBD, NTD and S2 subunits, like those observed in Fab fragment interactions of the above antibodies. We did not observe S1 shedding here, via Cryo-EM, likely due to the proline substitutions (6P) and a mutated furin cleavage site incorporated into our S-ectodomain construct to stabilize the prefusion conformation of the spike (36).

Binding of the spike trimer to the ACE2 receptor extends the RBDs outwards resulting in S1 shedding, which ultimately allows viral entry (8). The afore antibodies have been shown to act as ‘molecular ratchets’, biasing the SARS-CoV-2 spike conformational equilibrium toward opened RBDs, resulting in rotation beyond 20°, and hyperextension outwards to various extents, in order to accommodate Fab fragment binding. Subsequently, this causes the premature release of the S_1_ subunit. MAb S2A4 (PDB:7JVC) (31), wrenches the RBD, resulting in a 48° rotation and a 15 Å hyperextension, when compared to ACE2-bound spike-ectodomain (PDB:7A98) (8) (**Fig. S6**). For the 7F-RBD complex, we observe a 55° rotation of the RBD and an 8 Å hyperextension compared to the ACE2-bound spike-ectodomain (**Fig. S6**). The similarities in conformations we have found between the 7F-complex and previously published structures where S1 shedding is induced, suggest 7F exerts its antiviral activity through a similar mechanism.

### 7F does not interfere with the RBD-ACE2 interaction

To understand the mechanism of virus neutralization, we assessed the possible interference of 7F with S-mediated receptor-binding activity. An ELISA-based receptor binding inhibition assay revealed that 7F does not block the binding of SARS-CoV-2 spike ectodomain to the human ACE2 receptor (**Fig. 3A**). Structural alignment of an ACE2 RBD structure (PDB: 6VW1) (37) with a single molecule of 7F and single RBD showed that the interaction ‘major-interface’ would not hinder ACE2 access to the spike RBD (**Fig. 3B**). These results further suggest, the major interface interaction is the main driving force of 7F neutralization and trimer dimer formation may be a phenomenon observed at the higher concentrations of 7F and S-ecto used in our cryo-EM experiments.

**Figure 3.**
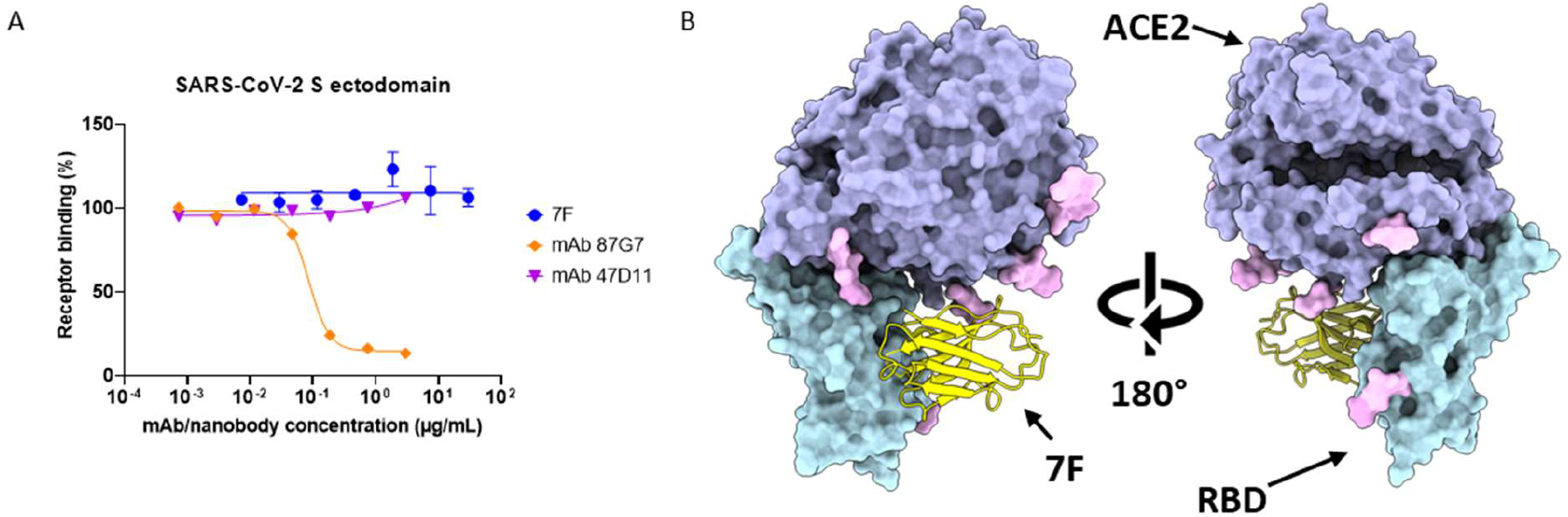
Functional and structural exploration of role of 7F in spike ACE2 interaction. **A.** ELISA-based receptor binding inhibition assay. SARS-CoV-2 spike ectodomain preincubated with serially diluted 7F or control mAbs 87G7 (ACE2 binding competitor) and 47D11 (not competing with ACE2 binding), were added to a plate coated with soluble human ACE2. The spike-ACE2 interaction was quantified using HRP-conjugated antibody targeting the C-terminal Strep-tag fused to SARS-CoV-2 spike ectodomain. Data points represent the mean ± SDM of n = 3 replicates from one representative of three independent experiments. Concentration displayed in µg/mL corresponds to range of 0.001 – 1000 nM. **B**. Superimposition of human ACE2/SARS-CoV-2 complex (PDB: 6VW1) locally refined model of SARS-CoV-2 RBD-7F complex protein structure.

### Cryo-EM reveals the interactions of 7F with the RBD

The major interface between 7F and the spike protein is characterized by a single salt bridge connecting D99 of 7F and K378 of RBD, along with eight hydrogen bonds (**Fig. 4**). The extensive interactions occur mainly through the long CDR3 loop of 7F, forming hydrogen bonds between D99, S102, F104, Y105, R108, E111, and consecutive residues of the RBD beta strand S371-T379. Additionally, two hydrogen bonds of the CDR2 loop, E57 and Y59, further stabilize the interaction with residues S383 and T385 of the same beta strand.

**Figure 4.**
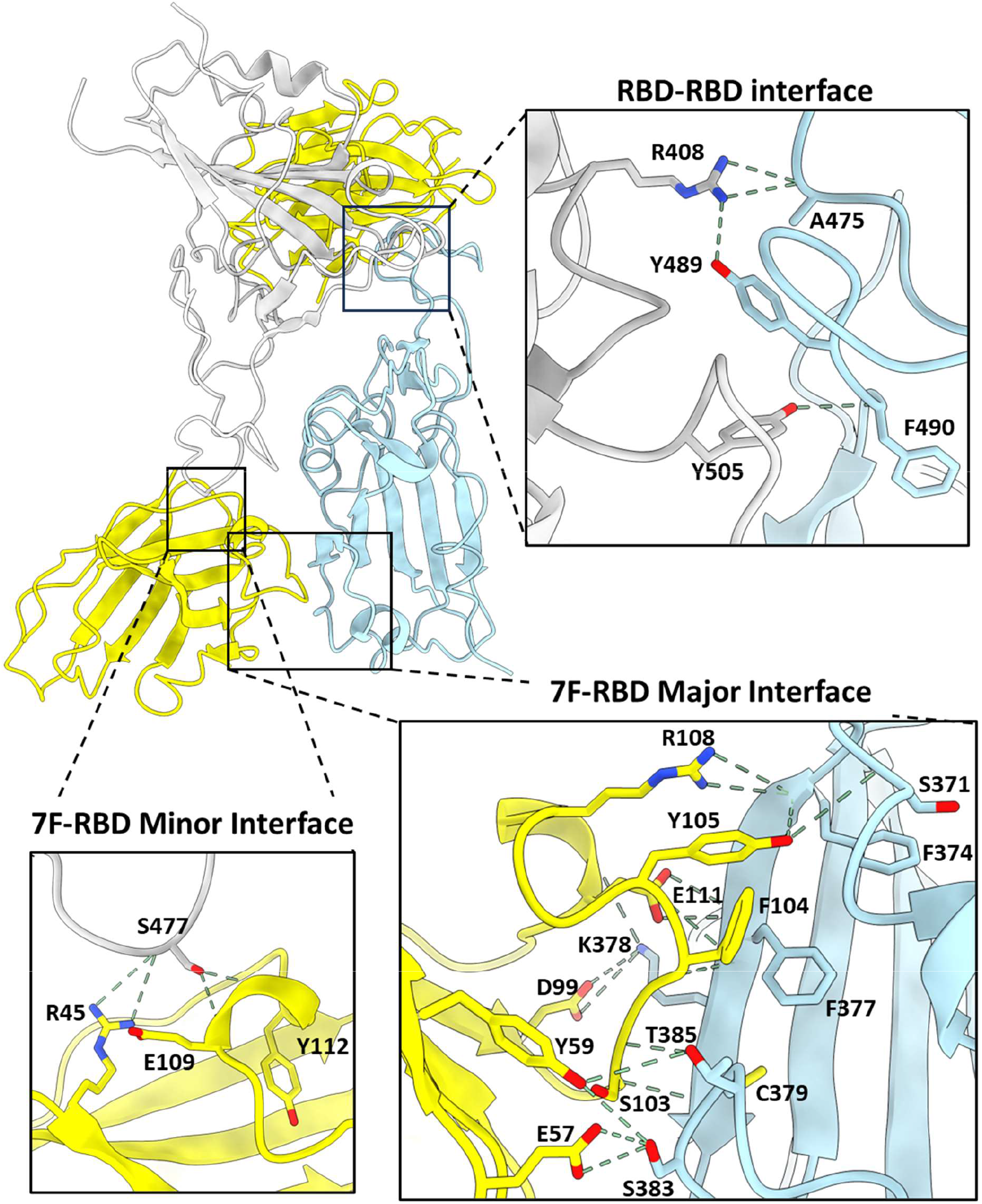
Structural basis for sdAb-7F binding to the SARS-CoV-2 RBD. Ribbon representation of the dimeric RBD-7F complex, with inset panels showing detailed interactions for the major and minor interfaces.

On the other hand, the minor interface involves only three hydrogen bonds, each binding to a single RBD amino acid S477 (**Fig. 4**). Interestingly, one of these interactions involves a residue, R45, from the framework of 7F, while the other two residues are from the CDR3 loop. The close interactions of both major and minor interfaces appear to facilitate the interaction between the RBD monomers, leading to extensive hydrogen bonding between G404, R408, G485, Y489, F490, and Y505 of one RBD with mirrored residues of the other RBD and resulting in a third interface, RBD-RBD.

While rare, trimer dimer formation has been induced by both mAb 6M6 (34) and nanobody Fu2 (24). Additionally, the SARS-CoV-2 Kappa variant formed head-to-head trimer dimers without the involvement of binding fragments (38). Although each of the binding molecules induces the formation of distinct trimer dimers, they share some aspects of the interfaces formed with 7F-RBD interactions described here (**Fig. 4**). The CDR3 of Fu2 similarly plays a predominant role in its major interface, forming a salt bridge with K378, and additional hydrogen bonds with T376-Y380 of the RBD beta strand, overlapping with residues reported here. Similarly, the formation of RBD-RBD interactions in trimer dimers formed by mAb 6M6 involves similar residues and interfaces as 7F. One notable difference between these molecules is that 7F-mediated trimer dimer formation is observed only with the trimers of SARS-CoV-2 adopting a three-up RBD formation, whereas 6M6 and Fu2 form these trimers with only two-up RBDs from different spike trimers of SARS-CoV-2.

### Dimerization of 7F enhances neutralization potency

Since the major-interface of 7F is a class IV epitope of the SARS-CoV-2 RBD - a region of the RBD known to be highly conserved amongst sarbecoviruses - we aimed to assess the conservation of the 7F binding site within the subgenus *Sarbecovirus* (28). For this, we mapped conserved and variable residues onto the RBD of the monomeric 7F-RBD complex (**Fig. 5A**). We found that all interacting residues of the RBD showed high conservation across selected sarbecoviruses, suggesting that 7F may exhibit binding and be able to neutralize many other sarbecoviruses and SARS-like CoVs, besides SARS-CoV-2, SARS-CoV and SARS-like CoV WIV16 tested here.

**Figure 5.**
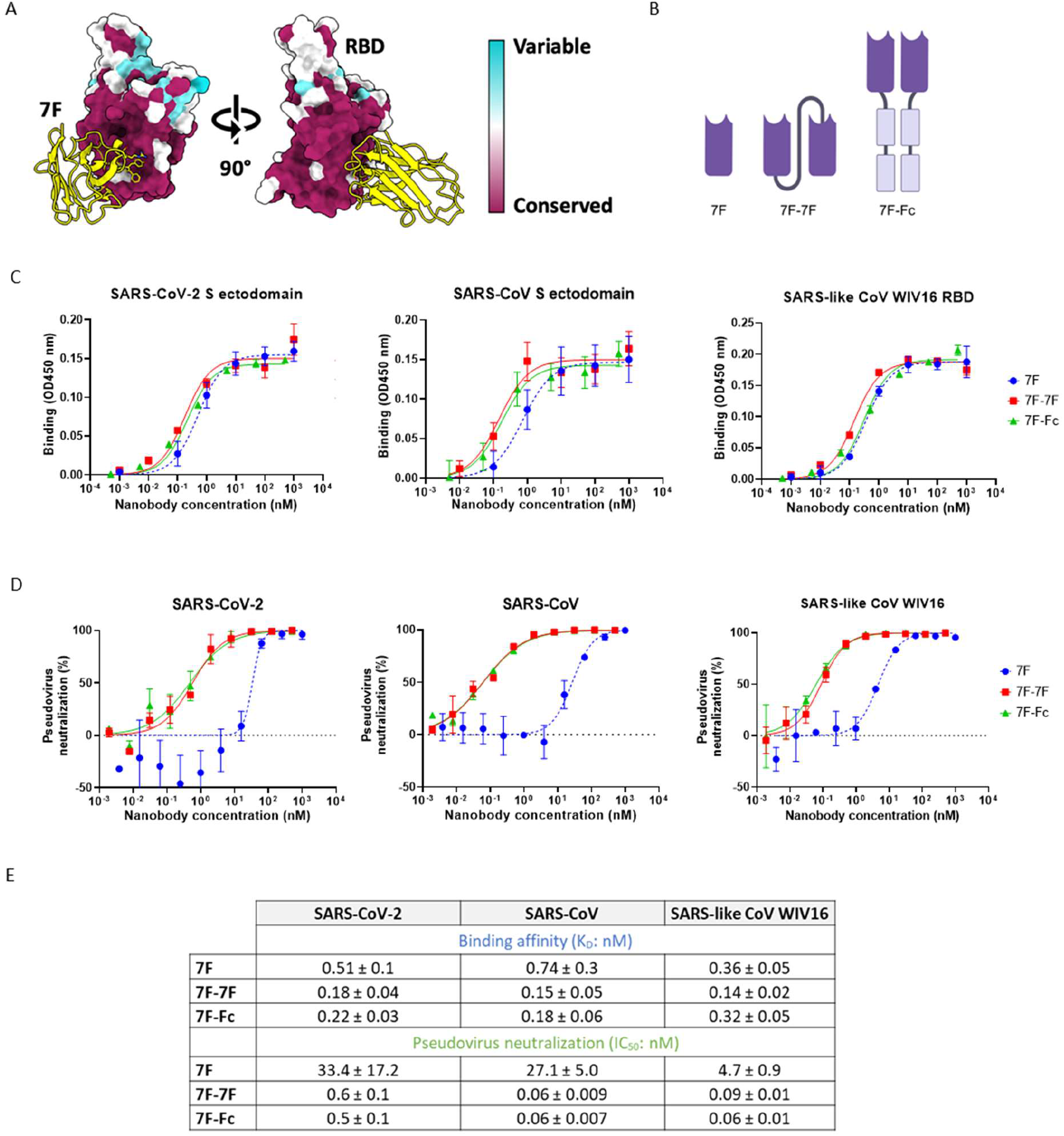
Enhanced neutralization potency of bivalent 7F constructs. **A.** Mapping of sarbecovirus amino acid conservation onto the surface representation of SARS-CoV-2 RBD in complex with 7F. **B.** Design of different 7F constructs, from left to right: 7F monomer, genetically linked bivalent 7F-7F and 7F-Fc fusion **C.** ELISA binding curves showing nanobody binding to immobilized SARS-CoV-2 spike ectodomain (left panel), SARS-CoV spike ectodomain (middle panel) and SARS-like CoV WIV16 RBD (right panel). Data points represent the mean ± SDM, for n = 3 replicates from one representative of three independent experiments **D**. Nanobody mediated neutralization of luciferase encoding VSV particles pseudotyped with spike proteins of (left panel to right panel) SARS-CoV-2, SARS-CoV and SARS-like CoV WIV16 on VeroE6 cells. Data points represent the mean ± SDM, for n = 3 replicates from one representative of two independent experiments **E.** Table showing the ELISA-based apparent binding affinities (K_D_) and pseudovirus neutralization based IC_50_ values of the monovalent and bivalent 7F constructs against SARS-CoV-2, SARS-CoV and SARS-like CoV WIV16 proteins or pseudoviruses, respectively. IC_50_ and K_D_ values (± standard deviation) were calculated from the binding and neutralization curves displayed in **B** and **C**, respectively.

Nanobody multimerization is known for its potential to enhance binding affinity, neutralization efficacy, and restore loss of binding and neutralization potency due to the antigenic drift in SARS-CoV-2, such as in [16,38]. Therefore, we engineered two distinct bivalent nanobody constructs. The first leverages insights from the cryo-EM structure, facilitating precise structure guided design of a genetically linked bivalent nanobody, 7F-7F. This construct features a glycine-serine linker of 10 amino acids long, that should enable simultaneous binding of two nanobodies to the spike trimer dimer complex. Complementing this structural approach, we developed a bivalent Fc-fused nanobody, 7F-Fc (**Fig. 5B**). Correct assembly of the Fc-fused construct by interchain disulfide bonds was confirmed by the higher molecular weight bands on the native gel, compared to the monomer based bands confirmed in the SDS gel (**Fig. S7**).

To ensure a fair comparison between monovalent and bivalent nanobodies, the constructs were evaluated based on their molar concentration, i.e. 7F (15 kDa), 7F-7F (30 kDa) and 7F-Fc (80 kDa). In the ELISA based binding assays, all 7F constructs showed high affinity binding, with apparent K_D_ values in the low nM range, to spike proteins of SARS-CoV and SARS-CoV-2 and RBD of SARS-like-CoV WIV16 (**Fig. 5C, E**). Despite these promising high binding affinities, only a very small benefit of the bivalent constructs over monovalent 7F was observed (**Fig. 5C**). The lack of substantial improvement in apparent binding affinities suggest this ELISA may not reflect the avidity effect expected to be observed for the bivalent nanobodies. Similar to the observations made with the monovalent nanobody, the bivalent 7F constructs exhibited no interference with the spike-ACE2 interaction (**Fig. S8**).

In contrast to the small change in apparent binding affinities, a substantial increase in neutralization potency was observed for both the 7F-7F and 7F-Fc against SARS-CoV-2, SARS-CoV and SARS-like CoV WIV16 pseudoviruses (**Fig. 5D**). The increase in potency observed for the bivalent nanobodies ranged from 50-450 times, depending on the strain tested (**Fig. 5E**). The enhanced neutralization observed was comparable for both 7F-7F and 7F-Fc, neutralizing the tested sarbecovirus pseudoviruses with IC_50_ values in the sub nM range.

### Dimerization of 7F increases neutralization breadth against SARS-CoV-2 variants of concern

Structural insight into 7F’s ability to inhibit SARS-CoV-2 variants of concern was gained by comparison of interacting spike residues between these variants and 7F (**Fig. 6A**). Within 7F’s major interface, we observed a single mutation, S471L in Omicron BA.1, which further evolved to S471F in later variants (**Fig. S9**). We anticipated that this mutation may abolish binding and/or neutralization of SARS-CoV-2 from Omicron and beyond. Consequently, we evaluated neutralization of our constructs against SARS-CoV-2 Omicron BA.2 and BA.5 pseudoviruses. As anticipated, neither VOCs were neutralized by monovalent 7F due to the 471 mutations. However, strikingly, in both bivalent forms, 7F recovered neutralization activity against both Omicron BA.2 and BA.5 (**Fig 6B**), with IC_50_ values in the low micromolar (µM) range (**Fig. 6C**). This suggests that 7F, in the bivalent format, can overcome both S471L and S471F mutations found in VOCs. Notably, both 7F-7F and 7F-Fc displayed comparable neutralization potencies against these VOCs. It is reasonable to assume that the bivalent nanobody constructs maintain neutralization potency against all VOCs included those most recent JN.1 and BA.2.86, as these variants have not accumulated further mutations in the 7F interacting residues. However, if the trimer dimer phenomenon does contribute to binding and neutralization of 7F, there are further mutations which have been acquired in later VOCs in both the minor-interface and RBD-RBD interface, which could hamper trimer dimer formation and therefore impact neutralization potency (**Table S1**).

**Figure 6.**
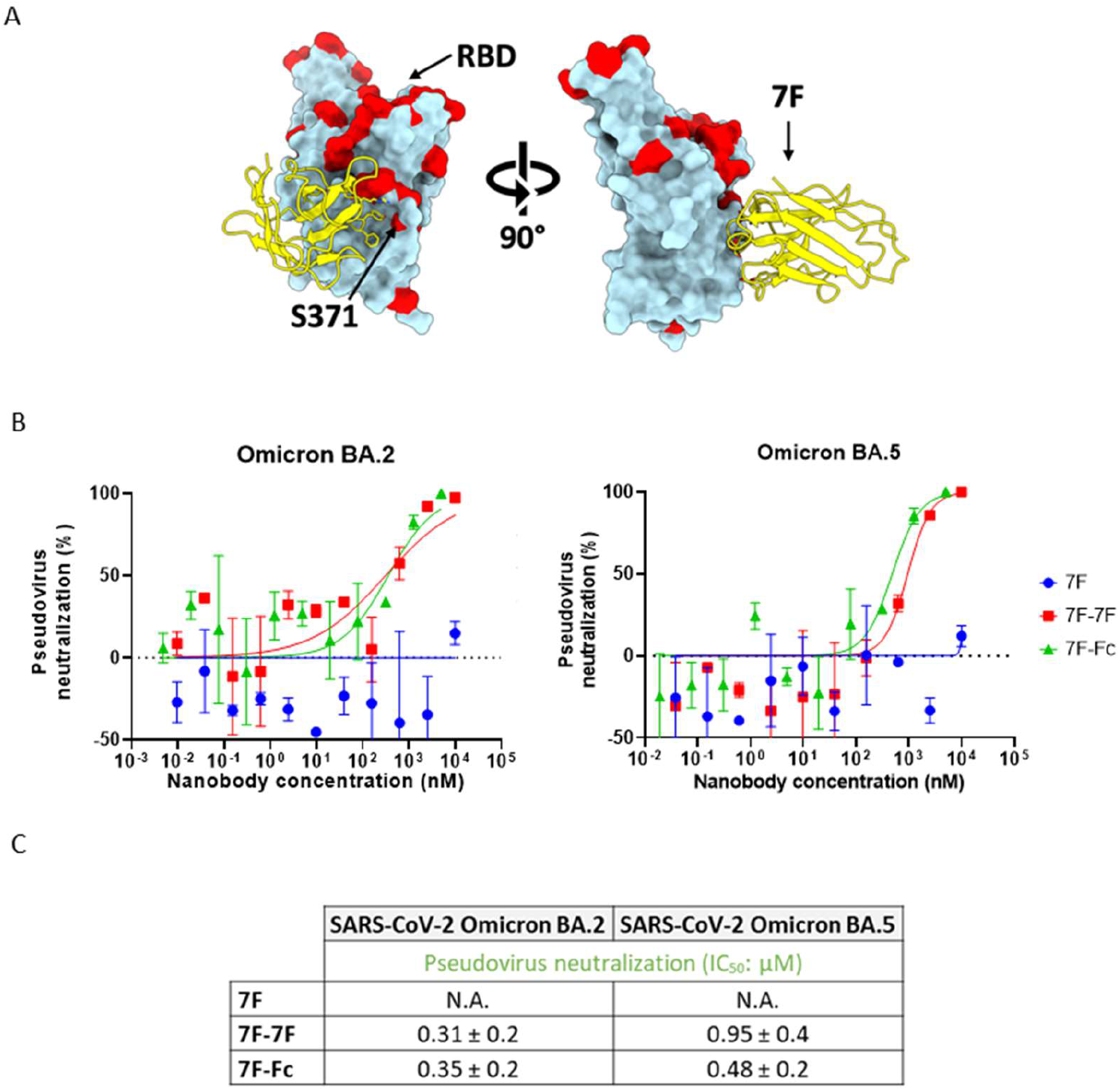
Dimerization increases the neutralization breadth of 7F against SARS-CoV-2 variants-of-concern. **A.** Mapping of SARS-CoV-2 amino acid conservation onto the surface of SARS-CoV-2 RBD in complex with 7F. **B.** Nanobody mediated neutralization of luciferase encoding VSV particles pseudotyped with spike proteins of (left) SARS-CoV-2 Omicron BA.2 and (right) SARS-CoV-2 Omicron BA.5 on VeroE6 cells. Data points represent the mean ± SDM, for n = 3 replicates from one representative of two independent experiments. **C.** Summary of the IC_50_ values of the monovalent and bivalent 7F constructs against SARS-CoV-2 Omicron BA.2 and BA.5 pseudoviruses. IC_50_ values (± standard deviation) were calculated from the neutralization curves displayed in **B**.

### 7F constructs effectively neutralize authentic SARS-CoV-2 infection in a fully differentiated 3D cell model of the human airway epithelium

We next investigated the neutralization potential of the 7F antibody formats against authentic SARS-CoV-2 infection in A549^ACE2+TMPRSS2+^ cells. Consistent with the observations for SARS-CoV-2 pseudovirus, the bivalent nanobodies demonstrate increased potency compared to their monovalent counterparts in neutralizing authentic SARS-CoV-2, with the IC_50_ value improving 3-5 fold (**Fig. 7A**). Furthermore, to explore the neutralization potency of the nanobodies in a more clinical relevant setting, we utilized human air-liquid interface (ALI) cultures of primary human airway epithelium (HAE) as an *ex vivo* model to replicate human infection. Our findings reveal that all three nanobody constructs exhibited a potent response against authentic SARS-CoV-2 in HAE culture, with 100 nM of nanobody being sufficient to completely inhibit infection over a 72 hour time period (**Fig. 7B**). Interestingly, in contrast to our observation in the virus neutralization assay, 100 nM nanobody surpassed the potency of 10 µM remdesivir against SARS-CoV-2 in the HAE culture.

**Figure 7.**
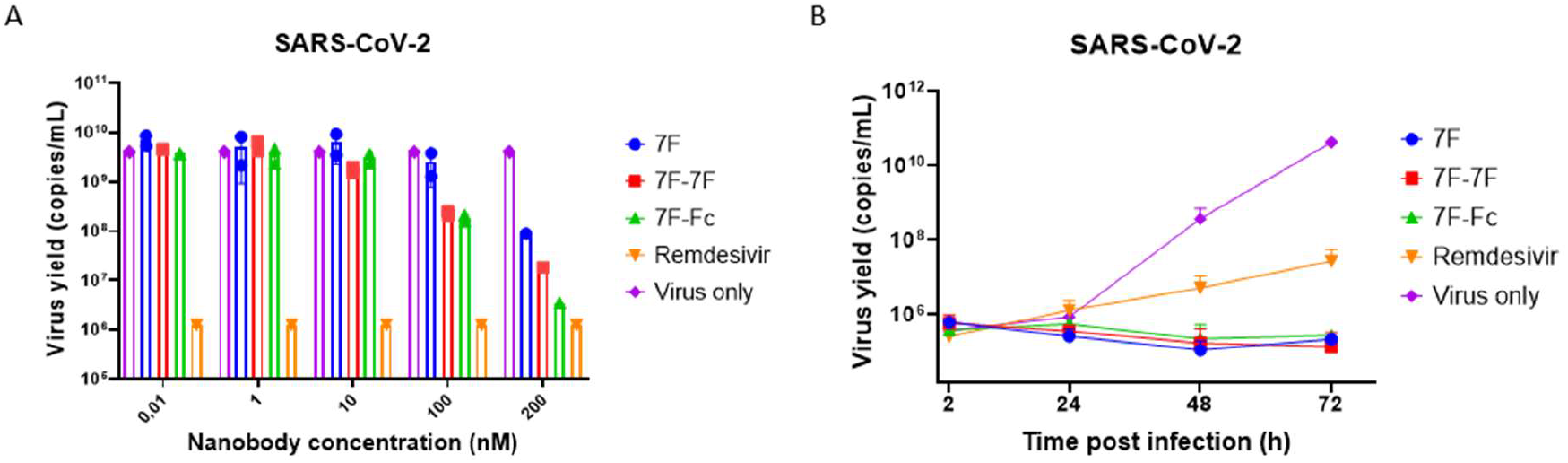
Monovalent and bivalent 7F constructs neutralize authentic SARS-CoV-2 in A549 cells and HAE cell cultures. **A.** Neutralization of SARS-CoV-2 in A549^ACE2+TMPRSS2+^ cells. Virus was pre-incubated with serial diluted nanobody, or 10 µM remdesivir, for 30 min before infecting A549^ACE2+TMPRSS2+^ cells. Infection was quantified by measuring the virus yield (viral RNA copies/ml, as determined with RT-qPCR) in cell culture supernatants of SARS-CoV-2 infected cells. **B.** Neutralization of SARS-CoV-2 in HAE cell culture. HAE cultures were incubated with SARS-CoV-2 and 100 nM nanobodies or 10 µM remdesivir on the apical side for 2 hours. Nanobody incubation was repeated every 24 hours. Graph showing the quantification of viral replication in the cultures, evaluated by RT-qPCR.

## Discussion

In this study, we present a broadly reactive nanobody, 7F, that is targeting a conserved site in the spike proteins of several sarbecoviruses. Through structural and functional analyses, we elucidated 7F’s ability to achieve cross-neutralization of SARS-CoV-2, SARS-CoV, and SARS-like CoV WIV16 pseudoviruses. 7F targets a conserved class IV epitope on the spike RBD, facilitating neutralization through a mechanism independent of receptor-binding inhibition. Our cryo-EM studies demonstrated that 7F induces the formation of spike trimer-dimers, mediated by interactions with both major and minor interfaces, on the spike RBD alongside a further RBD-RBD interface, likely induced by the stabilization of the RBDs in a head-to-head configuration. Interestingly, these interactions not only involve residues within the nanobody itself, but also highlight the role of RBD-RBD interactions in forming these spike trimer dimers.

Alongside the SARS-CoV-2 Kappa variant, capable of forming spike trimer dimers independently (38), CoV trimer-dimer formation has been reported previously for a nanobody and an antibody (24,34). The full structural basis was explored for one of these trimer dimer forming nanobodies, Fu2. Neutralization of Fu2 can partially be attributed to the aggregation of virions due to the trimer dimer formation (24). This mechanism may provide an explanation for the improved neutralization by 7F, though we could not validate this in our *in vitro* studies. While 7F-induced spike trimer-dimer formation does not appear to be the driver of neutralization, our structure does provide insights into nanobody induced higher order oligomerization, which could inform the design of new biparatopic molecules in the future. Unlike some other class IV nanobodies, 7F does not impede the spike-ACE2 interaction, suggesting an alternative mode of action (19,25,40). Possibly 7F achieves neutralization by destabilizing the trimeric spike structure, a mechanism previously observed with other class IV-targeting antibodies (32,35,40). ACE2 binding results in the release of S1 and ultimately viral entry. Here, we observed a structural similarity with comparable hyperextension and rotation of the RBD to accommodate 7F binding, features typically associated with class IV S1 shedding antibodies. However, we did not observe shedding in this instance, likely because of stabilizing mutations to the spike ectodomain to facilitate structural studies (32,35).

7F exhibits broad neutralization of SARS-CoV-2, SARS-CoV and SARS-like CoV WIV16 with IC50 values in the nM range, even in its monovalent form. Monovalent nanobodies displaying higher SARS-CoV-2 neutralization potency, sometimes in the picomolar range, have been described in literature, such as in [38,40]. However, these nanobodies only neutralize SARS-CoV-2 and therefore present a lower neutralization breadth compared to 7F. Additionally, potent broad neutralization of sarbecoviruses by other nanobodies has only been achieved after nanobody dimerization, as observed for VHH72 and Fu2 (24,25). Dimerization of 7F resulted in further improved potency of neutralization, with only a small change in binding affinity. Additionally, dimerization restored neutralization against SARS-CoV-2 VOCs, for which neutralization by monovalent 7F was lost with the S371L mutant in Omicron lineages and beyond. Given that S371L is the only mutation detected within the 7F epitope among the VOCs to date, we anticipate the bivalent 7F constructs to remain active against these variants. The neutralization breadth of 7F towards SARS-CoV-2 and other sarbecoviruses is a significant advantage and highlights its therapeutic potential to combat novel emerging sarbecoviruses.

The bivalent 7F-7F construct was specifically designed to facilitate the binding of two RBDs within one spike trimer simultaneously. However, it is possible that 7F also facilitates the crosslinking of spike trimers on the same or different viral particles (intra-virion and inter-virion cross-linking of spikes), potentially explaining the observed increase in neutralization potency in the absence of substantial increase in binding affinity. Interestingly, both 7F-7F and 7F-Fc exhibited equivalent potency in all conducted experiments. The option of generating bivalency through an Fc domain could further enhance protection *in vivo* through Fc-mediated antibody effector functions. While the bivalent 7F-7F could have an advantage in distribution *in vivo*, due to its smaller dimensions.

The use of HAE cell cultures in our study provides a physiologically relevant model for investigating respiratory infections. Comprising various epithelial cell types, HAE cultures exhibit a phenotypic resemblance to *in vivo* respiratory epithelium (42,43). Our findings highlight the efficacy of 7F in inhibiting viral infection and spread in this 3D human airway epithelial cell model. Even at low concentrations, 7F demonstrated potent neutralization of authentic SARS-CoV-2 infection in the HAE cell model, even surpassing the effectiveness of remdesivir. This promising result not only highlights the importance of using physiological relevant models in *in vitro* testing, but also supports the therapeutic potential of 7F and emphasizes the promise of this nanobody for further *in vivo* exploration.

As sarbecoviruses mainly affect the respiratory tracts, intranasal administration of 7F would be preferred. Previous studies have reported successful intranasal administration of genetically fused and Fc fused SARS-CoV-2 targeting nanobodies in mice and hamster models (18,44–46). Additionally, trimer dimer forming nanobody Fu2 showed potent protection of mice following intraperitoneal administration (24). These findings suggest that intranasal delivery of bivalent 7F could be a viable approach.

In conclusion, we have identified a broadly neutralizing nanobody, 7F, with anti-sarbecovirus activity. 7F was found to target a conserved class IV epitope on the spike RBD of sarbecoviruses, using a binding mechanism that results in the formation of spike trimer-dimers. Functional and structural data reveal the robust efficacy and molecular basis underlying the neutralization activity of 7F. With its potential to effectively combat infections caused by SARS-CoV, SARS-CoV-2, and related SARS-like coronaviruses, 7F emerges as a promising candidate for addressing future outbreaks of emerging and re-emerging sarbecovirus related diseases in humans.

## Materials and methods

### Nanobody selection

Two llamas were immunized with prefusion trimeric spike proteins of multiple betacoronaviruses; OC43, HKU1, MERS-CoV, SARS-CoV-1 and SARS-CoV-2. The animals received five immunizations in total, starting with SARS-CoV-2 spike followed by the other four spike proteins. Serum collected from the animals before and after immunizations was analyzed in ELISA to check for CoV spike reactivity. From blood collected on day 50, two phage display nanobody libraries (204 and 205) were constructed by an external company (QVQ BV). The libraries combined were used as input for nanobody panning to isolate sarbecovirus spike RBD targeting nanobodies. Two consecutive rounds of panning were performed, for the first round 0.5 µg/mL SARS-CoV-2-RBD protein was coated onto a NUNC Maxisorp strip (Thermo Fisher Scientific) at 4°C overnight. Coated wells were washed 3x using Phosphate-Buffered Saline (PBS) supplemented with 0.1% Tween-20 (PBS-T) and blocked using PBS supplemented with 4% w/v skimmed milk powder (PBSM-4%; Merck Millipore) for 1h at room temperature (RT). Antigen bound phages were eluted using 100 mM TEA buffer pH 12 and subsequently neutralized using 1M Tris-HCI pH 7.5. Eluted phages were amplified in exponentially growing *E.coli* TG1 cells which were infected with M13 helper phages. Phages were purified by a precipitation step using PEG-8000 and NaCl, and used as input for a second round of panning on 5 µg/mL SARS-CoV-2 RBD protein. Enrichment after selection was evaluated by titrating infected bacteria on Lysogeny broth (LB) agar plates containing 2% glucose and 100 µg/ml Ampicillin (amp). The output was tested in phage ELISA and periplasmic based ELISA. Promising clones, demonstrating binding to SARS-CoV-2 and SARS-CoV-1 S, were sequenced, and unique sequences of nanobodies were produced and tested again in ELISA for target specificity before being cloned into the pET21a expression vector.

### Constructing dimeric nanobodies

To obtain bivalent nanobody 7F-Fc, monovalent nanobody 7F was cloned into a pCG2 plasmid containing the human Fc-domain, using in-fusion cloning. PCR products were generated with overlapping primers (ordered from IDT) for the backbone and insert. Agarose gel electrophoresis was used to separate the backbone and the insert from each other, the appropriate bands were cut out and purified according to gel purification kit protocol (NucleoSpin). A 1:5 ligation, of purified backbone and insert, was performed using NEBuilder® HiFi DNA Assembly Master Mix (New England Biolabs). The ligation product was transformed into DH10B *E.coli* bacteria by heat shock. Success of the molecular cloning was confirmed with Sanger Sequencing performed by Macrogen Europe. The bivalent nanobody, 7F-7F, was constructed through the genetic fusion of two monovalent nanobody sequences connected by a 10 AA glycine serine linker.

### Production of nanobodies

Both monovalent 7F and bivalent 7F-7F were produced from a pET21a vector in *E.coli* BL21-DE3 cells. The culture was grown until OD_600_ of ∼1 after which cells were induced with 1 mM Isopropyl ß-D-1-thiogalactopyranoside (IPTG). After induction, temperature was reduced from to 25°C and the culture was incubated overnight. Nanobodies were retrieved from the periplasm by performing two freeze thaw steps and were purified via HisTrap HP columns (Thermo Fisher Scientific) and subsequent size-exclusion chromatography using the superdex 75 10/300 GL column (Cytiva) on the ÄKTAXpress chromatography system.

7F-Fc was expressed in Freestyle HEK293-F cells. Transfection of the plasmids into HEK293F was performed using polyethyleneimine (PEI) diluted in Opti-MEM (Gibco). After 24 hours, 300 mM Valporic acid (Sigma-Aldrich) and 10% primatone peptone (Sigma-Aldrich) was added to the transfected cells, which were then incubated until proteins could be harvested. Collected supernatant containing the Fc fused nanobody was incubated overnight with protein-A Sepharose beads (GE Healthcare), after which proteins were purified using poly-prep chromatography columns (Bio-Rad). Bivalent Fc nanobodies were eluted with 0,1 M citric acid (Sigma Aldrich) at pH 2,7. After elution, 3 M Tris-HCl (Sigma Aldrich) at pH 8,8 was added to neutralize the acidic elution. Nanobody concentrations were quantified by absorbance using the NanoDrop (Thermo Fisher Scientific). The purity of the nanobodies was determined by SDS-PAGE and Tris-Glycine gels.

### Cells

A549 cells (*Homo sapiens*, lung carcinoma, ATCC CCL-185) expressing human ACE2 receptor protein and TMPRSS2 activating protease (A549^ACE2+TMPRSS2+^) (47) and African green monkey kidney (VeroE6) cells were maintained in Dulbecco’s modified Eagle’s medium (DMEM) supplemented with 10% FBS, 1 mM sodium pyruvate (Gibco), nonessential amino acids (Lonza), penicillin (100 IU/ml), and streptomycin (100 IU/ml). A549^ACE2+TMPRSS2+^ cells were supplemented with blasticidin S (10 µg/mLl Gibco) and puromycin (0.5 µg/mL; Thermo Fisher Scientific) to maintain the expression of ACE2 and TMPRSS2 in the cells. Cells were cultured at 37°C in a humidified CO_2_ incubator. Cell lines tested negative for mycoplasma.

Human airway epithelial (HAE) cells were purchased commercially or isolated from conductive airways resected from transplant patients. The study was approved by the bioethical committee of the Medical University of Silesia in Katowice, Poland (approval no KNW/0022/KB1/17/10, dated 16 February 2010). Written consent was obtained from all patients. All experiments were performed in accordance with relevant guidelines and regulations. In brief, cells were mechanically detached from the tissue after protease treatment. Subsequently, cells were transferred onto permeable Transwell insert supports (ɸ = 6.5 mm) and cultured in bronchial epithelial growth medium (BEGM). After the cells reached full confluence, the apical medium was removed, and the basolateral medium was replaced with air-liquid interface (ALI) medium. Cells were cultured for 4-6 weeks to form fully differentiated, pseudostratified mucociliary epithelium as previously described (48,49).

### Viruses

Human codon-optimized genes, which encode the spike proteins of SARS-CoV, SARS-like CoV WIV16, and SARS-CoV-2 (Wuhan, Omicron BA.2, BA.5), were synthesized by GenScript. The generation of the variety of pseudo-typed VSVs followed earlier described procedures (50). In brief, HEK-293T cells were transfected with pCAGGS expression vectors containing the different spike proteins, each with a C-terminal cytoplasmic tail truncated by 28, 18 or 19 residues (SARS-CoV, SARS-CoV-2 and SARS-like WIV16 respectively) to enhance cell surface expression. Subsequently, cells were infected with VSV G-pseudo-typed VSVΔG carrying the firefly luciferase reporter gene 48 hours post-transfection. After 24 hours, the supernatant was collected, filtered, and the pseudo-typed VSVs were titrated on VeroE6 cells.

The SARS-CoV-2 live virus strains used in the study was the SARS-CoV-2 clinical isolate PL1455, which was isolated in-house (hCoV-19/Poland/PL_P18/2020, GISAID accession number: EPI_ISL_451979). Viral stocks were generated by infecting monolayers of VeroE6 cells (ATCC CCL-81). The cytopathic effect was evaluated after 3 days of infection and the cell supernatants were collected, aliquoted, and stored at −80°C. Control samples from mock-infected (noninfected) cells were prepared in the same manner. Virus yield (TCID50/ml) was assessed by titration on confluent A549^ACE2+TMPRSS2+^ using the Reed and Muench method (51).

### Expression and purification of spike proteins

Trimeric recombinant spike proteins of SARS-CoV-2 and SARS-CoV, and RBD protein of SARS-like CoV WIV16 were expressed and purified as described previously (52). In short, coronavirus spike ectodomain protein of SARS-CoV-2 was expressed transiently in HEK-293T cells with a C-terminal trimerization motif and Strep-tag using the pCAGGS expression plasmid. Similarly, a pCAGGS expression vector encoding RBD protein of SARS-like CoV WIV16 C-terminally tagged with a strep-tag was expressed in HEK-293T. Coronavirus spike ectodomain of SARS-CoV fused with a C-terminal trimerization motif, a thrombin cleavage site and a strep-tag purification tag were in-frame cloned into pMT\Bip\V5\His expression vector. The protein was expressed in HEK-293T. All recombinant proteins were affinity purified from the culture supernatant by streptactin beads (IBA) purification. Purity and integrity of all purified recombinant proteins was checked by Coomassie stained SDS-PAGE.

### ELISA analysis of nanobody binding to CoV antigens

Nanobodies were tested for binding to recombinant SARS-CoV-2 and SARS-CoV spike protein and to SARS-like CoV WIV-16 RBD protein. 100 ng of protein per well was coated overnight at 4°C onto 96-well NUNC MaxiSorp plates (Thermo Fisher Scientific). Coated plates were washed 3x using PBS and subsequently blocked using PBS supplemented with 2% bovine serum albumin (BSA; Fitzgerald) for 1 hour at RT. Nanobodies were allowed to bind to antigen coated plates at tenfold serial dilutions, starting from 5 µM diluted in 2% BSA PBS, at RT for 2 hour. Nanobody binding to spike was detected using 1:2000 rabbit anti-VHH (QVQ BV) for 1 hour RT followed by 1:2000 IRDye800 conjugated goat-anti-rabbit antibody (Li-COR Biosciences) or 1:1000 goat-anti-rabbit HRPO (Bio-Rad) for 1 hour at RT. Read-out of the IRDye800 secondary antibody, in mean fluorescence intensity 800 (MFI 800), was performed on an Odyssey near-infrared scanner (Li-COR Biosciences). HRP activity was measured at OD 450 nm using tetramethylbenzidine substrate (BioFX) using the BioSPX 800 TS Microplate reader (Tecan). In both cases, apparent binding affinity, K_D_, values were calculated by one-site specific non-linear regression on the binding curves (GraphPad Prism version 8.0.2).

### Biolayer interferometry

Binding analysis was performed using biolayer interferometry on the Octet (ForteBio) at 25°C. All reagents were diluted in PBS. First, 7F (5µM) was immobilized onto Ni-NTA biosensors (ForteBio) for 20 min. After a brief washing step, antigen binding was performed by incubating the biosensor with 5 µg of recombinant SARS-CoV-2 RBD or NTD protein for 30 min. Data was analyzed using Octet Data Analysis 9.0 (FortéBio).

### ELISA based Receptor-binding inhibition assay

Recombinant ACE2 protein was coated on 96-well NUNC Maxisorp plates (Thermo Fisher Scientific) at 100 ng per well overnight at 4°C. Plates were washed 3x using PBS containing 0.05% Tween-20 and blocked with PBSM-5% for 1 hour at RT. Recombinant strep-tagged SARS-CoV-2 spike or RBD protein (5 nM) was incubated with 4 fold serial diluted antibody or nanobody, starting from 20 nM or 2 µM respectively, at RT for 2 hours. The starting concentration for the antibodies used was based on their known activity range within this assay. After incubation, the mixture was added to the ACE2 coated plates and incubated for 2 hours at 4°C. Binding of spike or RBD protein to ACE2 was detected using 1:2000 HRP-conjugated anti-StrepMAb (IBA). HRP activity was measured at 450 nm using tetramethylbenzidine substrate (BioFX) and an ELISA plate reader (EL-808, BioTek).

### Pseudotyped virus neutralization assay

Nanobody neutralization was tested against the pseudotyped VSVs. Four-fold serial diluted nanobodies were preincubated with an equal volume of pseudotyped VSV at RT for 1 hour and then inoculated on confluently grown VeroE6 cells. After a 20 hour incubation at 37°C and 5% CO_2_, cells were washed once with PBS and subsequently lysed using Passive lysis buffer (Promega). The expression of firefly luciferase was measured on a Berthold Centro LB 960 plate luminometer using d-luciferin as a substrate (Promega). The percentage of neutralization was calculated as the ratio of the reduction in luciferase readout in the presence of nanobody normalized to luciferase readout in the absence of nanobody. Half-maximal inhibitory concentrations (IC_50_) values were determined using four-parameter logistic regression (GraphPad Prism 8.0.2.).

### Live virus neutralization assay

Serial nanobody dilutions were mixed with SARS-CoV-2 (1:1 v/v), incubated for 30 min at RT, and overlaid on confluent A549^ACE2+TMPRSS2+^ cells in a 96-well plate. The final virus titer was 1600 TCID50/ml. The cells were then incubated for 2 hours at 37°C in an atmosphere containing 5% CO_2_. The virus-infected cells in the absence of nanobodies were used as a positive control and mock-infected cells were considered a negative control. Remdesivir (10 µM; Gilead Sciences) was used as a reference. The unbound virus-nanobody complexes were removed by washing them twice with PBS, and nanobodies were reapplied on the cells. Cell culture supernatants were collected 3 days post-infection (p.i.) for RT-qPCR analysis. Experiments were carried out three times with each sample tested in duplicate.

HAE cultures were apically infected with a mixture of SARS-CoV-2 PL1455 and nanobodies similarly to the A549^ACE2+TMPRSS2+^ cells. After incubation and washing the cultures, nanobodies were administered to the apical side of the inserts, incubated for 15 min at 37°C and samples were collected for RT-qPCR analysis. The administration and collection of the samples were repeated every 24 hours until 72 h p.i. Experiments in HAE cultures were performed twice with each sample tested in duplicate.

The isolation of viral RNA from cell culture supernatants was performed automatically using the MagnifiQ 96 Pathogen instant kit (A&A Biotechnology) and the KingFisher Flex System (Thermo Fisher Scientific) according to the manufacturer’s protocol. Subsequently, viral RNA was reverse transcribed and quantified using GoTaq Probe 1-Step RT-qPCR System kit (Promega) in the presence of the specific SARS-CoV-2 probe (5′-6-FAM-ACT TCC TCA AGG AAC AAC ATT GCC A-BHQ-1-3′; 200 nM) and primers (Forward: 5′-CAC ATT GGC ACC CGC AAT C-3′, 600 nM; Reverse: 5′-GAG GAA CGA GAA GAG GCT TG-3′, 800 nM). Appropriate standards for the N gene of the virus were prepared to evaluate the number of viral RNA molecules in the samples. The reaction was performed in a thermal cycler (CFX Touch Real-Time PCR Detection System, Bio-Rad) with the heating scheme as follows: 15 min at 45°C, 2 min at 95°C, 40 cycles of 15 s at 95°C and 1 min at 56°C.

### Cryo-electron microscopy sample preparation and data collection

For the spike-7F complex, 2.5 μl 6P stabilized SARS-CoV-2 S-ectodomain, at a concentration of 28 μM (based on the molecular weight of the spike protomer) was combined with 0.1 μl of 140 µM 7F and incubated for ∼15 min at room temperature. Immediately before blotting and plunge freezing, 0.5 μl of 0.1% (w/v) fluorinated octyl maltoside (FOM) was added to the sample, resulting in a final FOM concentration of 0.01% (w/v). The sample solution (3 μl) was applied to glow-discharged (20 mAmp, 30 sec, Quorum GloQube) Quantifoil R1.2/1.3 grids (Quantifoil Micro Tools GmbH), blotted for 5 s using blot force 0 and plunge frozen into liquid ethane using Vitrobot Mark IV (Thermo Fisher Scientific). The data were collected on a Thermo ScientificTM KriosTM G4 Cryo Transmission Electron Microscope (Cryo-TEM) equipped with Selectris X Imaging Filter (Thermo Fisher Scientific) and Falcon 4i Direct Electron Detector (Thermo Fisher Scientific) operated in Electron-Event representation (EER) mode. In total, 1,948 movies were collected at a nominal magnification of 165,000×, corresponding to a calibrated pixel size of 0.73 Å/pix over a defocus range of −0.75 to −1.5 μm. A full list of data collection parameters can be found in Table S2.

### Single particle image processing

Data processing was performed using the CryoSPARC Software package (53). After patch-motion and CTF correction, particles were picked using a blob picker, extracted 4x binned and subjected to 2D classification. Following 2D classification, particles belonging to class averages that displayed high-resolution detail were selected for ab-initio reconstruction into six classes. Particles belonging to the spike trimer dimer complex classes were re-extracted 1.5x binned, resulting in a pixel size of 2.19 Å. The spike-7F particles were subjected to non-uniform refinement with D3 symmetry (54). At this point, the global resolution of the complex was 3.3 Å, however, the interfaces between spike and nanobody was resolved to a lower resolution. To improve the local resolution, particles in the final D3 global reconstruction were symmetry expanded, a custom mask encompassing one RBD and one nanobody involved in the head-to-head spike trimer dimer formation was used to carry out a cryoSPARC local refinement (BETA). This markedly improved local resolution, with the epitope resolved to a resolution of 3.1 Å, enabling sufficient confidence for modelling this epitope-paratope regions. For a more detailed processing methodology, see Fig.S2 and Fig.S3.

### Model building and refinement

UCSF Chimera (55) (version 1.15.0) and Coot (56) (version 0.9.6) were used for model building. The structure of the SARS-CoV-2 spike glycoprotein previously resolved (PDB ID 7R40) (57) and AlphaFold2 generated 7F nanobody (58,59) was used as a starting point for modelling of the spike-7F complex. Models were individually rigid body fitted into the density map using the UCSF Chimera “Fit in map” tool and then combined. The resulting model was then edited in Coot using the ‘real-space refinement, carbohydrate module (60) and ‘sphere refinement’ tool. To improve fitting, Namdinator (61) was utilised, using molecular dynamics flexible fitting of all models. Following this, iterative rounds of manual fitting in Coot and real space refinement in Phenix (62) were carried out to improve rotamer, bond angle and Ramachandran outliers. During refinement with Phenix, secondary structure and non-crystallographic symmetry restraints were imposed. The final model was validated in Phenix with MolProbity (63), EMRinger (64) and fitted glycans validated using Privateer (65,66).

### Structure analysis and visualization

Interacting residues of SARS-CoV-2 spike-7F epitopes were identified using PDBePISA (67) and LigPlot+ (68). Figures were generated using UCSF ChimeraX (69). Structural biology applications used in this project were compiled and configured by SBGrid (70).

## Declarations

## Ethics approval and consent to participate

Not applicable

## Consent for publication

Not applicable

## Availability of data and materials

All data generated or analyzed during this study are included in this published article [and its supplementary information files]. The cryo-EM maps and atomic structures have been deposited in the Protein Data Bank (PDB) Electron Microscopy Data Bank (EMDB) under accession codes: 9FR3 and EMD-50707 for SARS-CoV-2 S with nanobody-7F (global), and 9FR4 and EMD-50708 for SARS-CoV-2 S with nanobody-7F (local). Source data are provided with this paper.

## Author Contributions

ICS, OJD, DLH, BJB, SO designed the research project; ICS, OJD, planned and executed experiments; ID cryo-EM grid preparation and data collection; OJD and DLH cryo-EM data processing; OJD model building; TB, MC, MB assisted with experiments; AZ, KP executed and reviewed live virus and HAE culture experiment; ICS, OJD analyzed data and wrote the manuscript; ICS, OJD made the figures; DLH, BJB, SO reviewed data, and reviewed and edited the manuscript; CAMH, SO acquired the funding. All authors reviewed the manuscript.

## Acknowledgements

Not applicable.

