## Supplementary information for "A Spike Trimer Dimer-Inducing Nanobody with Anti-Sarbecovirus Activity"

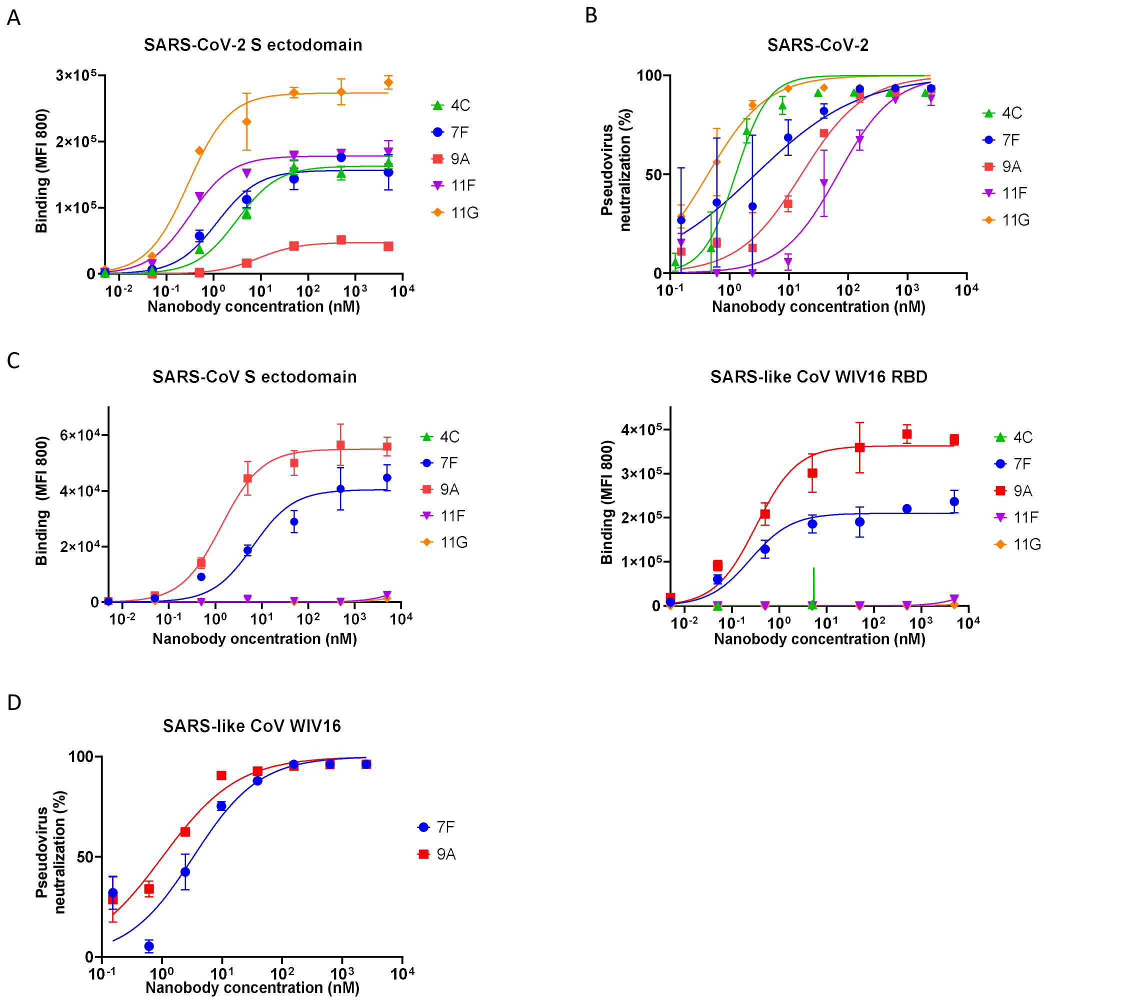


***Supplementary 1 Initial characterization of selected SARS-CoV-2 targeting nanobodies A.*** *ELISA binding curves showing nanobody binding to immobilized SARS-CoV-2 S ectodomain* ***B.*** *Nanobody mediated neutralization of luciferase encoding VSV particles pseudotyped with spike proteins of SARS-CoV-2 on VeroE6 cells.* ***C.*** *ELISA binding curves showing nanobody binding to immobilized SARS-CoV S ectodomain and SARS-like CoV WIV16 domain.* ***D.*** *Nanobody mediated neutralization of luciferase encoding VSV particles pseudotyped with spike proteins of SARS-like CoV WIV16 on VeroE6 cells. ELISA binding curves displayed in* ***A*** *and* ***C*** *show data points which represent the mean ± SDM of n = 3 replicates from one representative of three independent experiments. The pseudovirus neutralization graphs in* ***B*** *and* ***D*** *show data points which represent the mean ± SDM of n = 3 replicates from one representative of three independent experiments.*

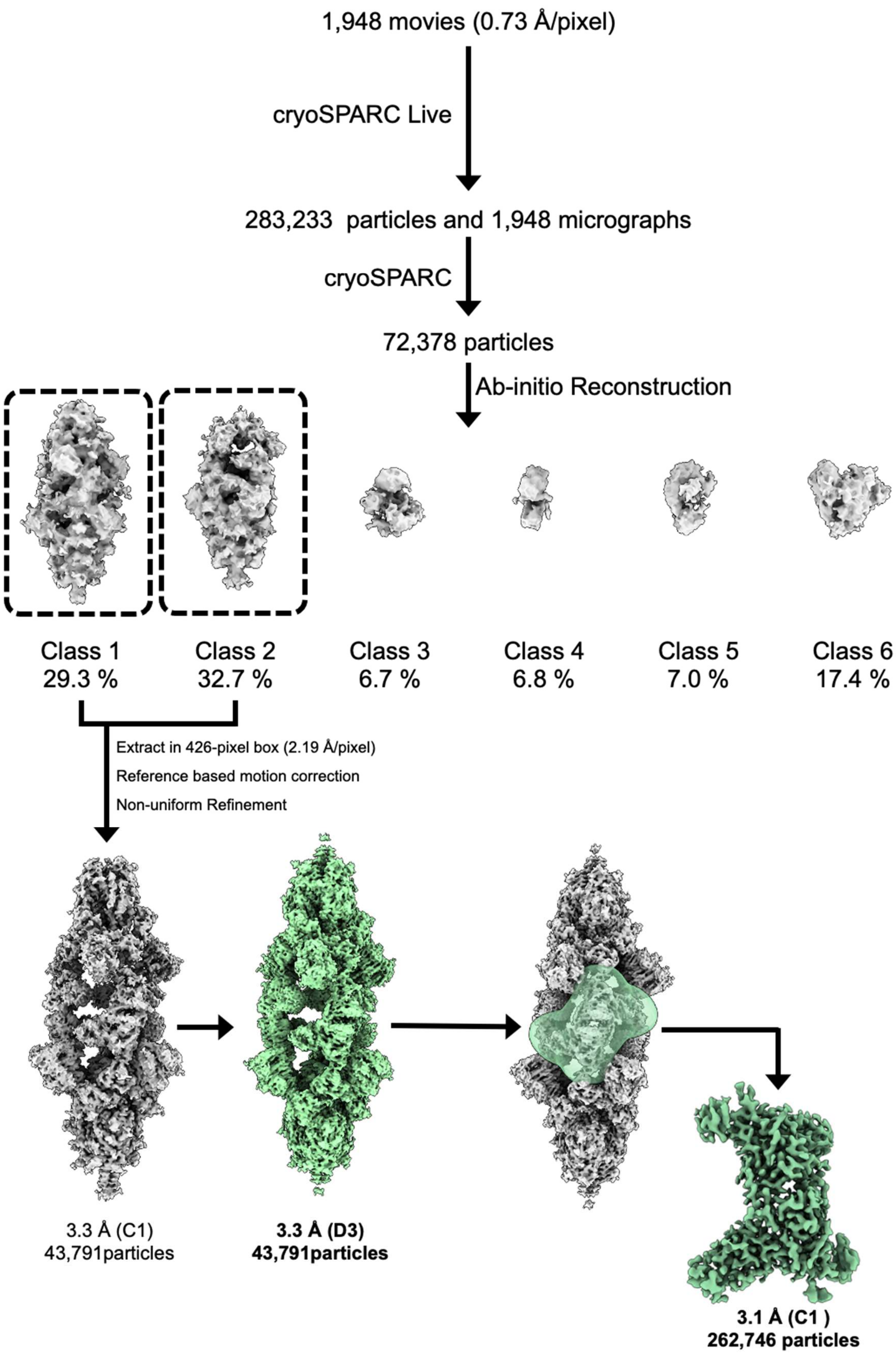


***Supplementary 2 Single-particle cryo-EM data processing pipeline for the SARS-CoV-2 S-7F nanobody complex.***

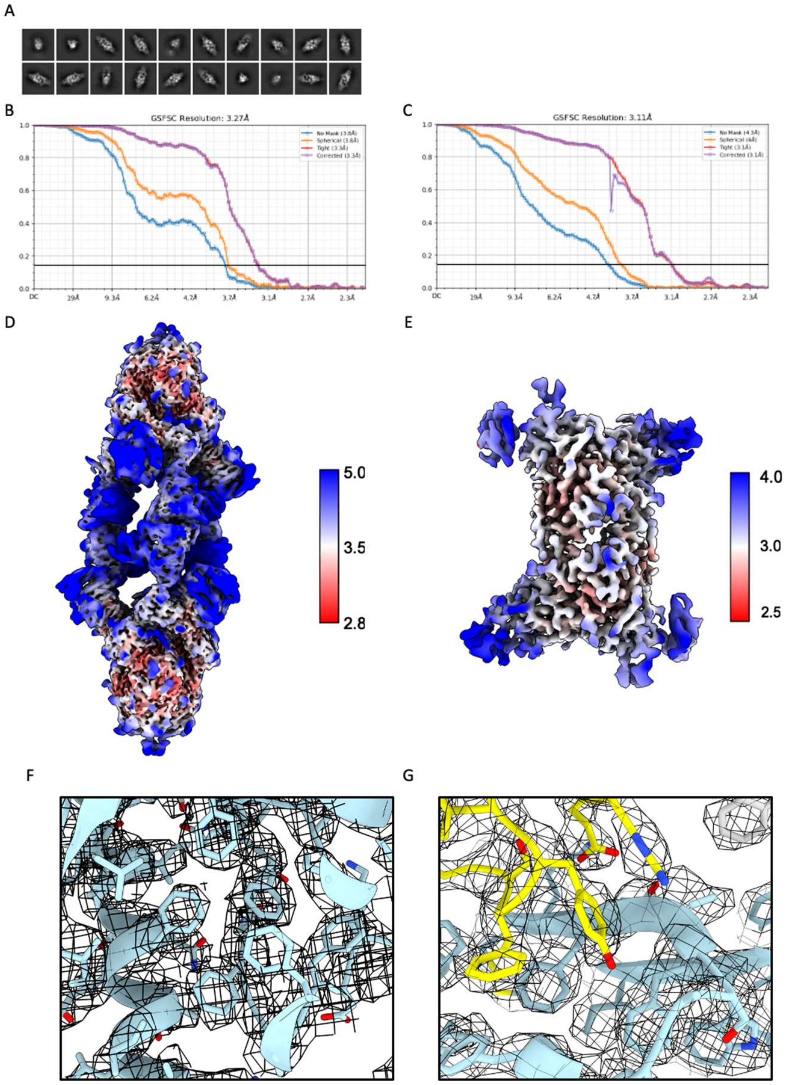


***Supplementary 3 Single-particle cryo-EM data processing for the SARS-CoV-2 S-7F nanobody complex. (A)*** *Selected 2D classes* ***(B)*** *Gold-standard Fourier shell correlation (FSC) curve generated from the independent half maps contributing to the 3.3 Å global resolution density map of the the SARS-CoV-2 spike in complex with 7F nanobody.* ***(C)*** *Gold-standard Fourier shell correlation (FSC) curve generated from the independent half maps contributing to the 3.1 Å local refined density map of the the SARS-CoV-2 spike-7F complex paratope-epitope.* ***(D)*** *Local resolution filtered EM density map for the D3 refined the SARS-CoV-2 Spike-7F complex, colored according to local resolution which was calculated in CryoSPARC.* ***(E)*** *Local resolution filtered EM density map for the local refinement of the SARS-CoV-2 Spike-7F complex paratope-epitope, colored according to local resolution which was calculated in CryoSPARC.* ***(F)****Representative view of the SARS-CoV-2 spike ectodomain fitted model in the EM density of 3.3 Å global resolution density map of the the SARS-CoV-2 spike in complex with 7F nanobody shown as a black mesh* ***(G)*** *Zoomed-in view of the interacting region of 7F and SARS-CoV-2 RBD with the EM density of the 3.1 Å local refined density map of the the SARS-CoV-2 Spike-7F complex paratope-epitope shown as a black mesh.*

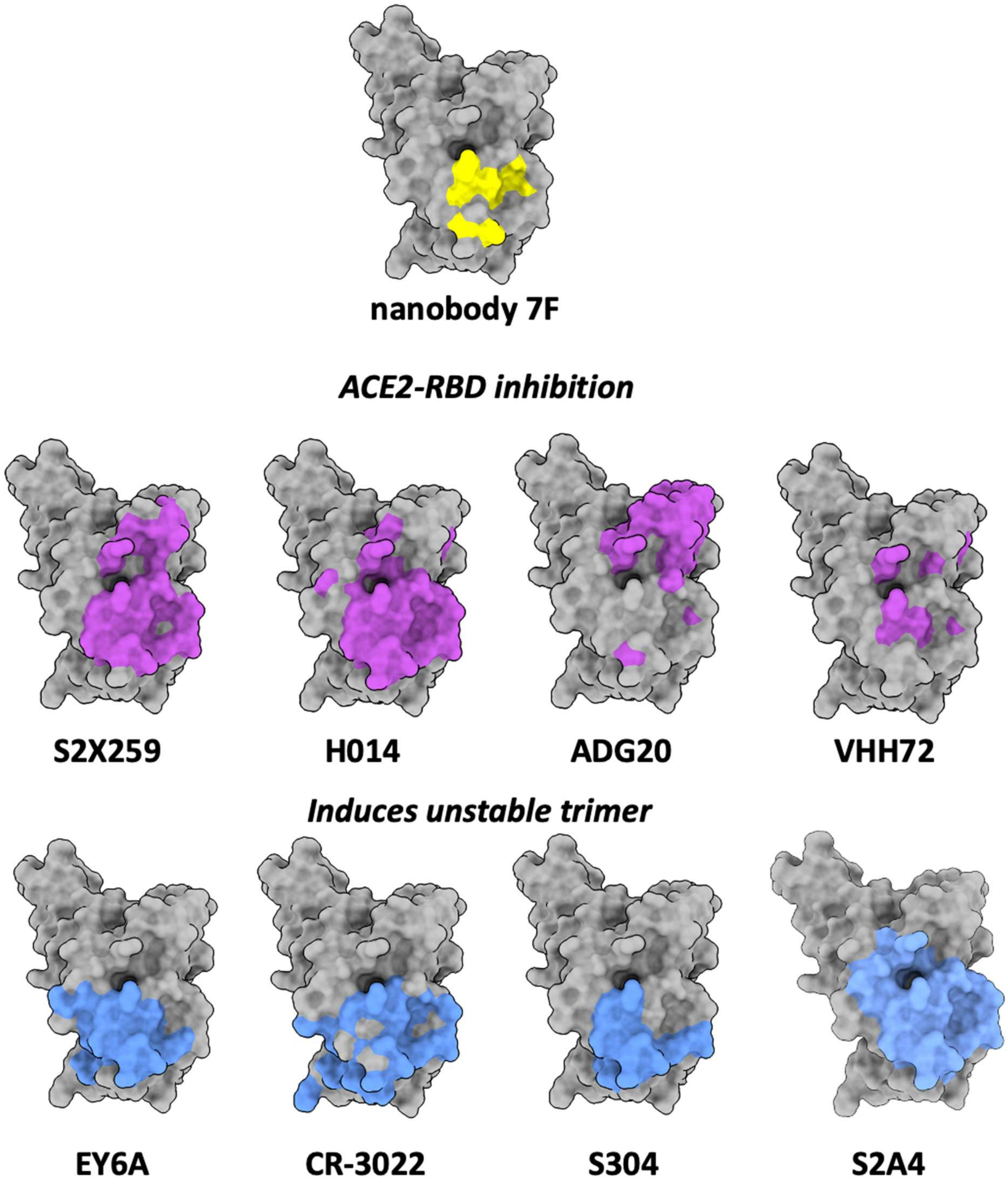


***Supplementary 4 The interacting site of 7F in comparison with other antibodies and nanobodies**. The interactions of 7F compared with S2X259 (PDB:7RAL), H014 (PDB:7CAH), ADG20 (PDB:7U2D), EY6A (6ZFO), CR-3022 (PDB: 8FAH), S304 (PDB: 7JW0), n3130v (PDB:8I4G), S2A4 (PDB:7JVA) and nanobody VHH72(PDB:6WAQ).*

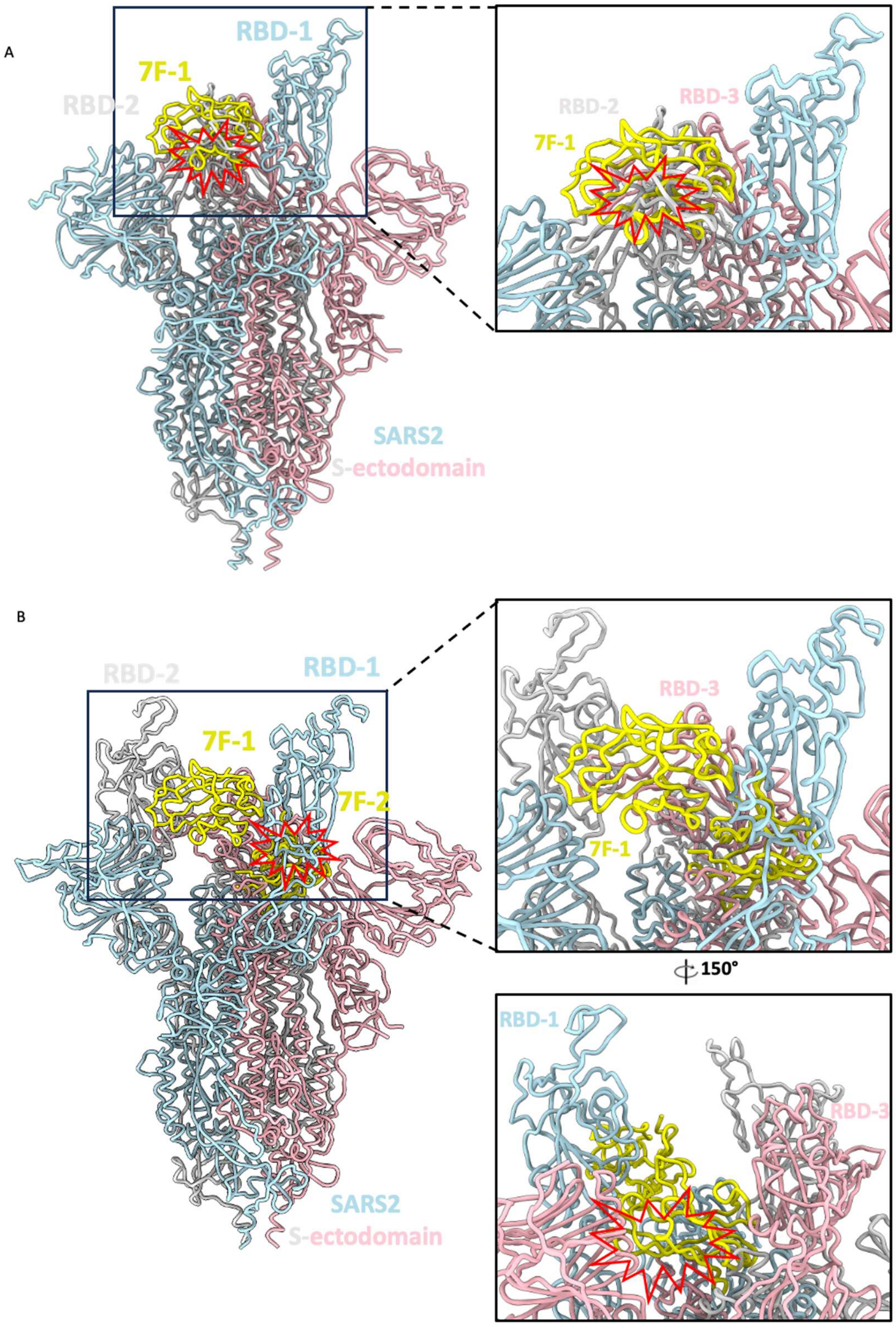


***Supplementary 5 Structural representation of 7F binding inaccessibility.*** *(A) SARS-CoV-2 S-ectodomain with "1 up" RBD monomer (PDB: 7VHN), with SARS-CoV-2 Spike-7F complex superimposed onto the open RBD (RBD-1). (B) SARS-CoV-2 S-ectodomain with "2 up" RBD monomer (PDB: 7A93), with two SARS-CoV-2 Spike-7F complex's superimposed onto one open RBD (RBD-1) and one closed RBD (RBD-3). Clashes which may induce steric hinderance are indicated with a red clash sign.*

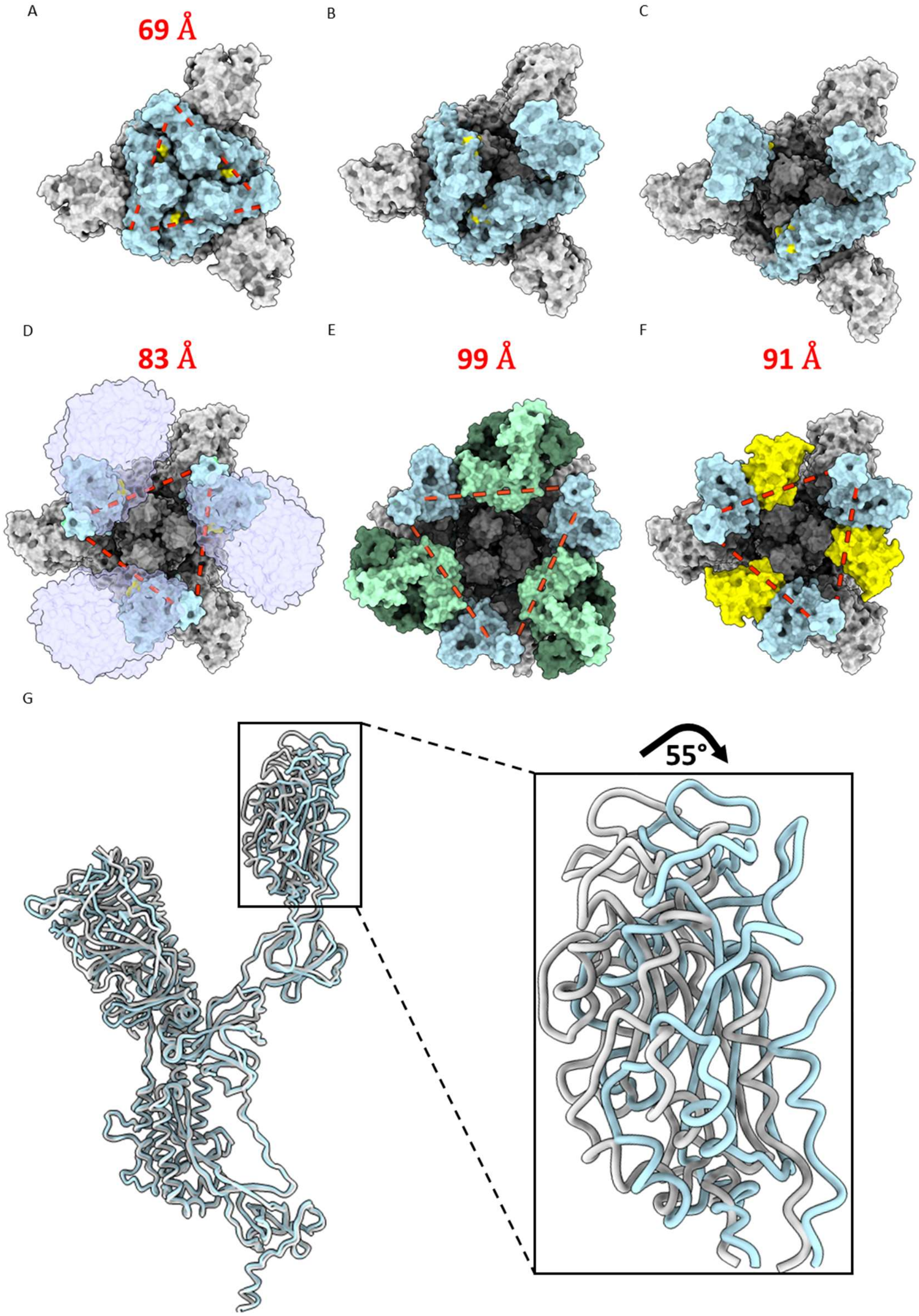


***Supplementary 6 Structural representation of potential S1 shedding mechanism induced by 7F binding.*** *(A-F) top views of SARS-CoV-2 S-ectodomain adopting: closed conformation (PDB:6XR8), "1 up" (PDB: 7VHN), "2 up" (PDB: 7A93), "3-up" in complex with ACE2 (PDB:7A98), "3-up" in complex with S2A4 (PDB:7JVC) and "3-up" in complex with 7F. Distances between each RBD indicated by red dashed lines are measured using ChimeraX and displayed in red for panel A, D, E and F. All S-ectodomains are represented in light grey, with RBDs in light blue. S2A4 Fab fragments are represented in green and 7F or 7F contact points are indicated in yellow. ACE2 is*

*represented in transparent lilac. (G) Superimposed SARS-CoV-2 S-protomers in complex with either ACE2 (light grey) or in complex with 7F (light blue). Angular hyperextension of the RBD was calculated in ChimeraX and represented.*

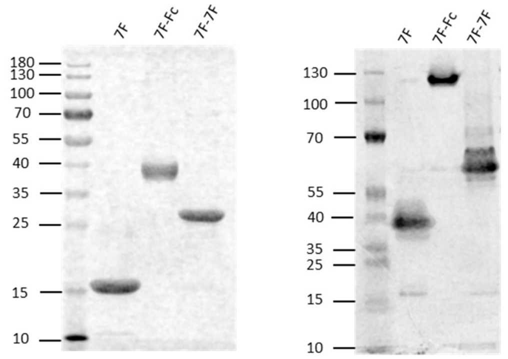


***Supplementary 7 Gel analysis proves correct formation of bivalent 7F nanobodies.*** *Nanobody integrity analysis by SDS-PAGE under reducing conditions (left) and native, non-reducing, tris-glycine gel (right), followed by Coomassie staining. The molecular weight marker does not accurately reflect protein migration in native PAGE due to differences in protein charge, size, and shape under native conditions.*

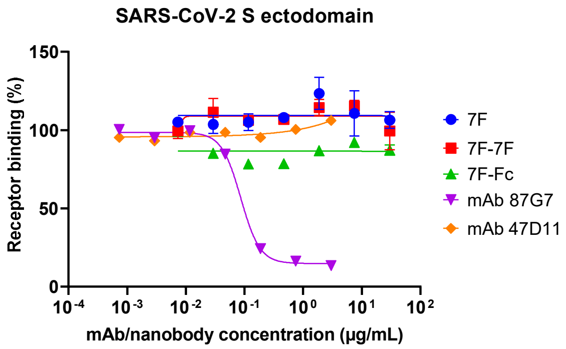


***Supplementary 8 Both monovalent and bivalent 7F constructs do not interfere with the RBD-ACE2 interaction.*** *ELISA-based receptor binding inhibition assay. SARS-CoV-2 spike ectodomain preincubated with serially diluted monovalent and bivalent 7F constructs or two control mAbs 87G7 (ACE2 binding competitor) and 47D11 (non ACE2 binder), was added to a plate coated with soluble human ACE2. The spike-ACE2 interaction was quantified using HRP-conjugated antibody targeting the C-terminal Strep-tag fused to SARS-CoV-2 spike ectodomain. Data points represent the average ± SDM, for n = 3 replicates from one representative of three independent experiments. Concentration displayed in µg/mL corresponds to range of 0.001 – 1000 nM.*

| | RBD-7F interface interacting residues | | | | | | | RBD-RBD interface interacting residues | | | | |
|---|---|---|---|---|---|---|---|---|---|---|---|---|
| | 371 | 374 | 377 | 379 | 383 | 385 | 477 | 408 | 475 | 489 | 490 | 505 |
| SARS-CoV-2 | | | | | | | | | | | | |
| Wuhan | S | F | F | C | S | T | S | R | A | Y | F | Y |
| Alpha | S | F | F | C | S | T | S | R | A | Y | F | Y |
| Beta | S | F | F | C | S | T | S | R | A | Y | F | Y |
| Gamma | S | F | F | C | S | T | S | R | A | Y | F | Y |
| Delta | S | F | F | C | S | T | S | R | A | Y | F | Y |
| Omicron BA.1 | L | F | F | C | S | T | N | R | A | Y | F | H |
| Omicron BA1.1 | L | F | F | C | S | T | N | R | A | Y | F | H |
| Omicron BA.2 | F | F | F | C | S | T | N | S | A | Y | F | H |
| Omicron BA.3 | F | F | F | C | S | T | N | S | A | Y | F | H |
| Omicron BA.4 | F | F | F | C | S | T | N | S | A | Y | F | H |
| Omicron BA.5 | F | F | F | C | S | T | N | S | A | Y | F | H |
| Omicron BQ1.1 | F | F | F | C | S | T | N | S | A | Y | F | H |
| Omicron XBB.1 | F | F | F | C | S | T | N | S | A | Y | S | H |
| Omicron XBB.1.5 | F | F | F | C | S | T | N | S | A | Y | S | H |
| Omicron XBB.1.9 | F | F | F | C | S | T | N | S | A | Y | S | H |
| Omicron EG.5.1 | F | F | F | C | S | T | N | S | A | Y | S | H |
| Omicron BA.2.86 | F | F | F | C | S | T | N | S | A | Y | F | H |
| Omicron JN.1 | F | F | F | C | S | T | N | S | A | Y | F | H |
| Omicron JN.1.4 | F | F | F | C | S | T | N | S | A | Y | F | H |
| Sarbecoviruses | | | | | | | | | | | | |
| SARS-CoV | S | F | F | C | S | T | G | R | P | Y | W | Y |
| SARS-like CoV WIV16 | S | F | F | C | S | T | G | R | P | Y | W | Y |
| RaTG13 | S | F | F | C | S | T | S | R | A | Y | Y | H |
| Merbecovirus | | | | | | | | | | | | |
| MERS-CoV | L | V | F | C | S | A | Y | S | N | V | W | V |

***Supplementary Table 1 Sequence conservation of the 7F-RBD and RBD-RBD interacting residues****. Aligned amino acid sequences of the residues involved in RBD-7F (left part of the table) or RBD-RBD (right part of the table) interactions, of SARS-CoV-2 variants of concern, a few sarbecoviruses and a merbecovirus. Dotted lines highlight the viruses tested in this study. Residues that differ from the SARS-CoV-2 Wuhan sequence are indicated In Orange.*

### Cryo-EM data collection, refinement and validation statistics

| | SARS-CoV-2-7F (Global) (EMDB-50707) (PDB 9FR3) | SARS-CoV-2-7F (Local) (EMDB-50708) (PDB 9FR4) |
|---|---|---|
| **Data collection and processing** | | |
| Magnification | 105,000 | 105,000 |
| Voltage (kV) | 300 | 300 |
| Electron exposure (e–/Å$^2$) | 50 | 50 |
| Defocus range (μm) | -1.2 to -2.0 | -1.2 to -2.0 |
| Pixel size (Å) | 0.73 | 0.73 |
| Symmetry imposed | D3 | C1 |
| Initial particle images (no.) | 282,223 | 282,223 |
| Final particle images (no.) | 43,791 | 262,746 |
| Map resolution (Å) | 3.3 | 3.1 |
| FSC threshold | 0.143 | 0.143 |
| Map resolution range (Å) | 0.0-33.8 | 0.0-49.2 |
| **Refinement** | | |
| Initial model used (PDB code) | 7R40 | 7R40 |
| Model resolution (Å) | 2.9 | 2.9 |
| FSC threshold | 0.143 | 0.143 |
| Map sharpening *B* factor (Å$^2$) | 62.0 | 121.4 |
| Model composition | | |
| Non-hydrogen atoms | 51756 | 4874 |
| Protein residues | 6630 | 614 |
| Ligands | 42 | 0 |
| *B* factors (Å$^2$) | | - |
| Protein | 48.79 | 47.45 |
| Ligand | 78.37 | - |
| R.m.s. deviations | | |
| Bond lengths (Å) | 0.006 | 0.004 |
| Bond angles (°) | 0.679 | 0.678 |
| Validation | | - |
| MolProbity score | 2.2 | 1.72 |
| Clashscore | 20.20 | 5.94 |
| Poor rotamers (%) | 0.00 | 0.00 |
| Ramachandran plot | | - |
| Favored (%) | 94.07 | 94.06 |
| Allowed (%) | 5.93 | 5.94 |
| Disallowed (%) | 0.00 | 0.00 |

*Supplementary table 2 Cryo-EM model building statistics.*
